# scDual-Seq of *Toxoplasma gondii*-infected mouse bone marrow-derived dendritic cells reveals host cell heterogeneity and differential infection dynamics

**DOI:** 10.1101/2023.01.13.523919

**Authors:** Franziska Hildebrandt, Mubasher Mohammed, Alexis Dziedziech, Amol Bhandage, Anna-Maria Divne, Fredrik Barrenäs, Antonio Barragan, Johan Henriksson, Johan Ankarklev

## Abstract

Host-parasite interactions include complex interplays between an invading and a defending organism, each continuously adapting to gain the upper hand. The protozoan parasite, *Toxoplasma gondii*, can invade every nucleated cell type in a vertebrate host, including immune cells while the host mounts a protective response.

Here, we utilize Dual-scSeq to parse out heterogeneous transcription of bone marrow-derived dendritic cells (BMDCs) infected with *T. gondii* type I, RH (LDM) or type II, ME49 (PTG) parasites, over multiple time points post infection.

We find that the two parasite lineages distinctly manipulate two subpopulations of infected BMDCs. Co-expression networks establish host and parasite genes, with implications for modulation of host immunity and host-pathogen interactions. Integration of published data validates immune pathways and suggests novel candidate genes involved in host-pathogen interactions. This study aims to provide a comprehensive resource for future characterization of host-pathogen interplay among other protozoan parasites within their host niches, as well as that of bacterial and viral pathogens.

## INTRODUCTION

The dynamics of infection are determined by interactions between the infectious agent, often invading microorganisms such as viruses, bacteria and parasites and their respective host. Infected host cells are required to recognize and respond to an intruder by activation of the immune system. Simultaneously, invading microorganisms ensure their survival and replication by evading the host’s immune response. Many obligate intracellular microorganisms are found to alter the cellular expression programs of infected and surrounding cells to actively evade immunity, scavenge nutrients and establish chronic infection or dormancy states within the host. Conversely, host cells have developed numerous strategies to prevent establishment of different pathogens including protective mechanisms that are specific for the intruder ^1^. Cell responses can be unique, not only based on the kind of pathogen but also based on which host cell is responding. Despite host challenges, intracellular parasites are highly successful in establishing infection evidenced by their global prevalence. However, knowledge on the dynamics of host-pathogen interactions remains fragmented.

The mammalian immune system heavily relies on innate immune sensors important in microbial perception ^2^. Two cell types of importance are the dendritic cells (DCs) and macrophages, which both present antigens to adaptive immune cells and can both be derived from bone-marrow in a laboratory setting. DCs and macrophages are some of the first cells to respond to infection as part of the mononuclear phagocyte system ^3^. DCs have developed very specific mechanisms to elicit sophisticated and precise immune responses. Macrophages on the other hand, are effector cells known for their role in phagocytosis of microbes ^4^.

Increasingly distinct subtypes of bone marrow-derived DCs (BMDCs) have been described, including cells which display profiles more similar to macrophages. These subtypes were also shown to display specific functions regarding particular pathogens or diseases ^3,5,6^. For example, LPS was found to elicit a bimodal transcriptional response in two subpopulations of BMDCs ^5,6^. The extent of heterogeneity within BMDC and other immune cell populations in their steady-state and in response to pathogens has yet to be explored extensively at the single-cell level. *Toxoplasma gondii*, known to invade and extensively modulate its host cells, deserves further exploration.

*Toxoplasma gondii* belongs to the Apicomplexan phylum including more than 4500 species of obligate intracellular, parasitic protozoa ^7^. Uniquely, *T. gondii* demonstrates the highest prevalence of infection where one third of the human population is estimated to be infected ^8^. Infection occurs by ingestion of undercooked meat containing parasite cysts or via consumption of contaminated food or water containing oocysts originating from feline feces ^9,10^. The majority of individuals infected with *Toxoplasma* are asymptomatic. However, vulnerable individuals including the immunocompromised, pregnant women and children born from an infected mother, can experience severe or even lethal disease in the form of disseminated or latent infections or cerebral toxoplasmosis ^10^.

An active infection is typically characterized by rapidly replicating tachyzoites, able to invade any nucleated host cell. The interaction between parasite-secreted proteins from organelles such as rhoptries, micronemes and dense granules within the apical complex and host adhesion receptors aid in the invasion process ^11,12^. Extensive and continuous replication of intracellular tachyzoites eventually leads to the disruption of the host cell. Thereafter parasites invade surrounding cells or disseminate to other areas of the body, even crossing barriers into the placenta or the brain ^13^.

DCs and macrophages, which are enriched in the intestinal lamina propria and the Peyer’s patches are the first responding immune cells towards incoming *Toxoplasma* parasites ^14^. The parasite can modulate the migratory behavior ^15–18^ or the immune response of infected Macrophages and DCs and eventually establish chronic infection through the formation of bradyzoites in the brain ^19^.

Across Europe and North America, the predominant infecting clonal lineages are commonly separated into type I, type II and type III parasites ^20^. These clonal lineages show differential correlations with disease progression and outcome in both mice and humans ^21,22^. Type I parasites, as opposed to type II or type III parasites, cause lethal virulence in mice ^22^. In contrast, type II parasites disseminate more effectively due to increased hypermotility and longer migratory distances in infected leukocytes, possibly giving them an advantage in the development of chronic infection ^23^. Further, clonal lineages display differences in macrophage activation, with type II parasites activating classical immune activation, while type I and type III parasites induce alternative macrophage activation ^24^.

Previous characterization of transcriptional changes upon infection with *T. gondii* was integral to gain insight in the host-response. However, current publicly available data is mostly limited to population-wide bulk RNA sequencing ^25,26^ of the host or in tandem with the parasite ^27^ as well as single cell RNA sequencing data of the host alone ^28^. Single cell dual sequencing (sc-DualSeq) enables dissecting and deciphering molecular population heterogeneity during highly dynamic interactions of an active infection ^29,30^. The applicability of sc-DualSeq and the relevance of heterogeneity in infection dynamics was recently demonstrated in *T. gondii* infected monocytes ^31^ and human foreskin fibroblasts ^32^. However, an exhaustive image of host-pathogen interactions between different *T. gondii* clonal lineages and murine leukocytes on the transcriptional level and in single cells over other relevant time points has yet to be described.

With heterogeneity present in infection at the single cell level, further exploration is required to understand the parallel transcriptional dynamics of the host and parasite over the course of infection. For example, through the use of scDual-Seq, orchestration of immune responses towards the parasite and identification of distinct cells whose behavior might deviate from the majority can be distinguished.

In this study, we aim to parse out the heterogeneity of infection kinetics, using scDual-Seq to explore the host-parasite interaction between murine BMDCs infected with types I and II *T. gondii*, to identify inter-species co-expression networks.

We uncover underlying factors of heterogeneity in the host cell population beyond inherent stochasticity of gene expression, including; hidden cell subtypes, cell cycle genetic markers and effects of parasite replication. Additionally, we aim to create a computational resource to compare transcriptional responses in host-pathogen complexes. The ambition is to enable the broader infection and immunology community to explore similarities between other host-parasite interactions, including but not limited to other protozoan pathogens, such as *Plasmodium spp, Cryptosporidum spp*, and *Leishmania spp*. Finally, our observations provide information with future clinical implications such as the development of effector molecule-targeting therapies, by providing novel candidate genes identified both in the host and parasite.

## RESULTS

### *T. gondii* infects two sub-populations of bone marrow-derived dendritic cells (BMDCs)

scDual-Seq enables the simultaneous recovery of host and pathogen signatures from individual BMDCs infected with two different strains of *T. gondii* parasites: *T. gondii* PTG and *T. gondii* LDM at 3 and 12 hours post infection (hpi). Between these time points, the parasites undergo at least one cycle of endodyogeny, a specialized replication mechanism of *T. gondii* tachyzoites ^33^, leading to two or four parasites within each host cell at 12 hpi. As parasite lysate is unable to actively invade and further modulate a host cell we only collected data for all controls at 3 hpi (figure 1a).

**Figure 1.**
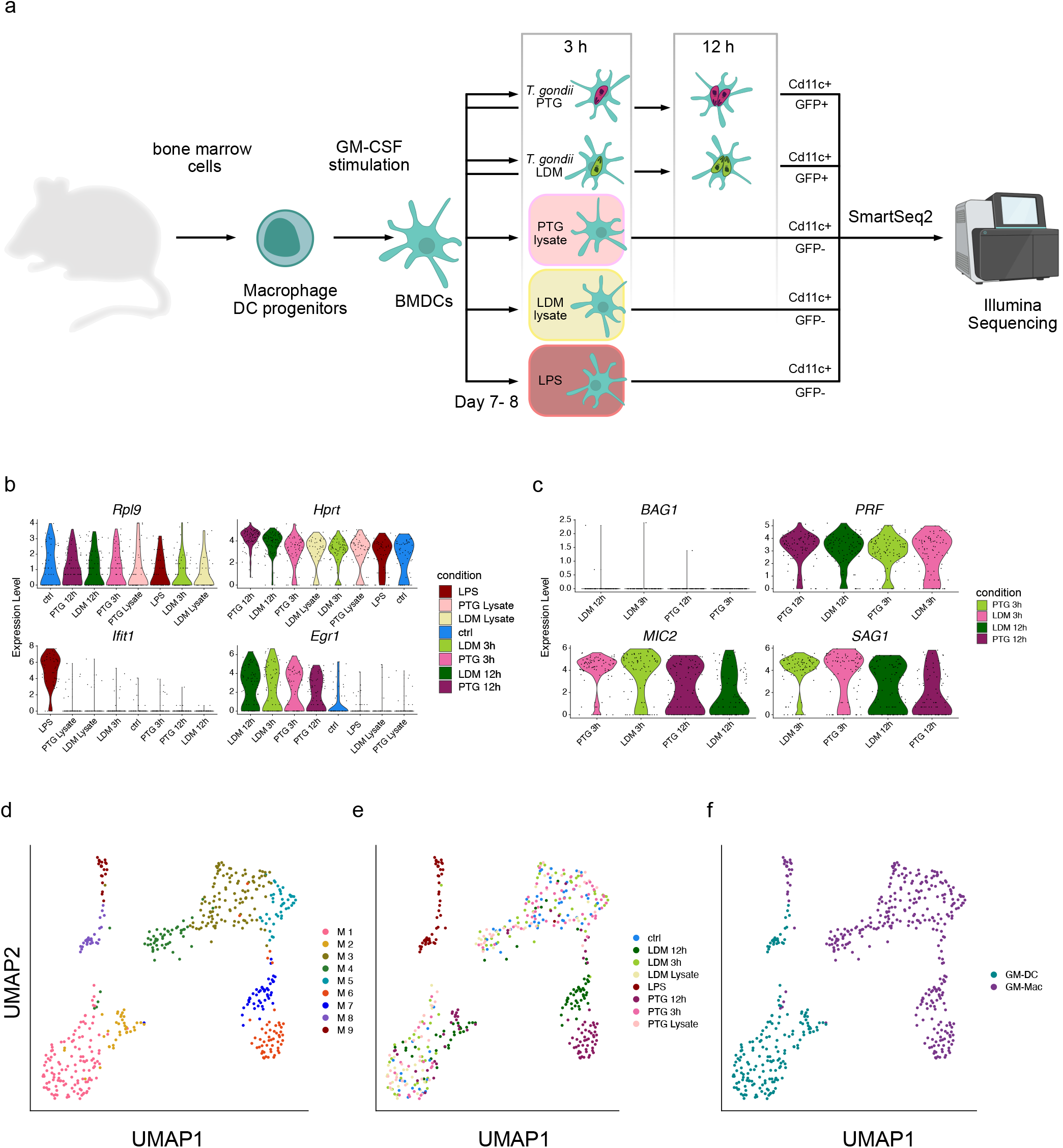
Transcriptional analysis of Toxoplasma BMDC infection at the single cell level. **a)** Schematic overview of study design and biological significance. Primary BMDCs were collected, activated and infected with *T. gondii* LDM (LDM) and PTG (PTG) for 3 and 12 hours. Heat-inactivated lysates and LPS served as controls. Cells were then sorted based on GFP+ fluorescence from the parasite and presence of the CD11C-PE-Cy7 surface marker and subjected to scRNA-seq and subsequent analysis. **b)** Distribution of expression levels across infection conditions of selected mouse marker genes ordered by expression levels from highest to lowest. **c** Distribution of selected *T. gondii* markers across original conditions ordered by expression levels from highest to lowest. **d)** UMAP projection of single cells colored by identified clusters from graph-based clustering. **e)** UMAP projection of single cells colored by infection condition. **f** UMAP projection of single cells colored by infected host cell type.

The majority of reads originated from host transcripts across all cells and *T. gondii* PTG transcripts decrease between 3 and 12 hpi, opposed to an increase of transcripts for LDM parasites. This indicates time-dependent differences in general transcriptional activity between strains (Supplementary figure 1).

To validate our approach, we inspected the distribution of established host- and parasite marker genes across conditions. Common housekeeping genes *Rpl9* and *Hprt* ^34,35^ show equal expression levels across all conditions and cells. *Ifit1* is a marker for immune response to LPS infection in macrophages ^36^ and is upregulated in LPS-stimulated cells but not in *T. gondii* infected cells. Conversely, only cells infected with *T. gondii* sustain previously reported high levels of *Egr1* expression ^37^, independent of their clonal lineage (LDM or PTG) (figure 1b).

The expression of the *T. gondii* marker gene, *BAG1*, exclusively expressed in bradyzoite stages ^38^, showed no upregulation in our data. *PRF*, essential for gliding motility, invasion and exit from host cells ^39^, shows increased expression in the late stages of *T. gondii* tachyzoites. *MIC2* and *SAG1*, two other *T. gondii* genes expressing proteins essential for invasion, virulence and host cell binding ^40,41^, showed expression across all infection conditions, peaking at early time points (figure 1c). The observed expression of established marker genes of *T. gondii* infection as well as expression profiles across positive (LPS) and negative controls (uninfected control and lysate controls) suggests that the number of cells in our study is sufficient to investigate host-pathogen interactions on the transcriptional level.

Next, we isolated expression data from both the host and the parasite. Unsupervised clustering of the host data resulted in 9 clusters (M1-M9) across all conditions (figure 1d). When we compared the cluster annotations of cells to the experimental conditions in a UMAP embedding, we observed that cluster M8 and M9 were comprised of only LPS activated cells, while cluster M2 and M6-M7 were comprised of almost exclusively *T. gondii* infected cells from 12 hpi which included both strains (LDM and PTG). Interestingly, we found that the distribution of cells from 3 hpi, uninfected cells and cells challenged with parasite lysate were uniform across the remaining clusters (M1 and M3-M5) (figure 1d,e, Supplementary figure 2).

Further, we observed four clearly separated groups of cells in the UMAP projection, including clusters M1-M2, M3-M5, M6-M7 and M8-M9, suggesting clear transcriptional differences between these cells (figure 1d). Single cell studies have previously enabled the investigation of bimodality in the expression of seemingly homogenous BMDC populations ^5,6^. The study performed by Helft and colleagues suggests that murine bone marrow cells cultivated with granulocyte-macrophage colony-stimulating factor (GM-CSF), results in a heterogeneous population of CD11c+ MHCII+ macrophages (GM-Macs) and dendritic cells (GM-DCs). Based on these observations, we sought to investigate whether we can distinguish heterogeneous cell types within our data. With Helft *et al*. as our reference, we were able to classify clusters M1-M2 and cluster M8 as GM-DCs and the remaining murine clusters M3-M7 and M9 as being predominantly composed of GM-Macs (figure 1f, supplementary figure 3, supplementary data 1). The majority of late stage infected (12 hpi) cells can be divided into three major clusters: I) M2, containing a mixture of LDM and PTG infected GM-DCs, II) M6, containing exclusively PTG infected GM-Macs and III) M7, containing exclusively LDM infected GM-Macs (figure 1e,f,). This indicates that different clonal lineages of *T. gondii* elicit different host responses in GM-Macs but not in GM-DCs at 12 hpi *in vitro*.

### Different clonal lineages of *T. gondii* evoke differential host responses in a cell type dependent manner

Parsing out the different transcriptional host responses of the cell subtypes across the 3 cluster groups highlighted above, we first performed a Differential Gene Expression Analysis (DGEA) between clusters M6 (PTG at 12 hpi) and M7 (LDM at 12 hpi). We then compared differentially expressed marker genes between the 3 clusters. Interestingly M2 cells had a transcriptional profile more similar to cluster M7 (figure 2a). Overall, we observed opposite transcriptional regulation of genes involved in innate immunity and inflammation between M6 (PTG at 12 hpi) and M7 (LDM at 12 hpi). Genes upregulated in M6 are associated with the GO-terms “response to bacterium” (GO:0009617) and “response to interferon-beta” (GO:0035456) (figure 2a, supplementary data 2). These genes are involved in the formation of complex signal transduction networks that activate innate immunity and inflammation ^42^ by the induction of immune mediators ^43^, Pattern Recognition Receptors (PRRs) ^44^, the JAK/STAT pathway ^45^, NF-κB signaling ^46^, and inflammasome activation ^47^.

**Figure 2.**
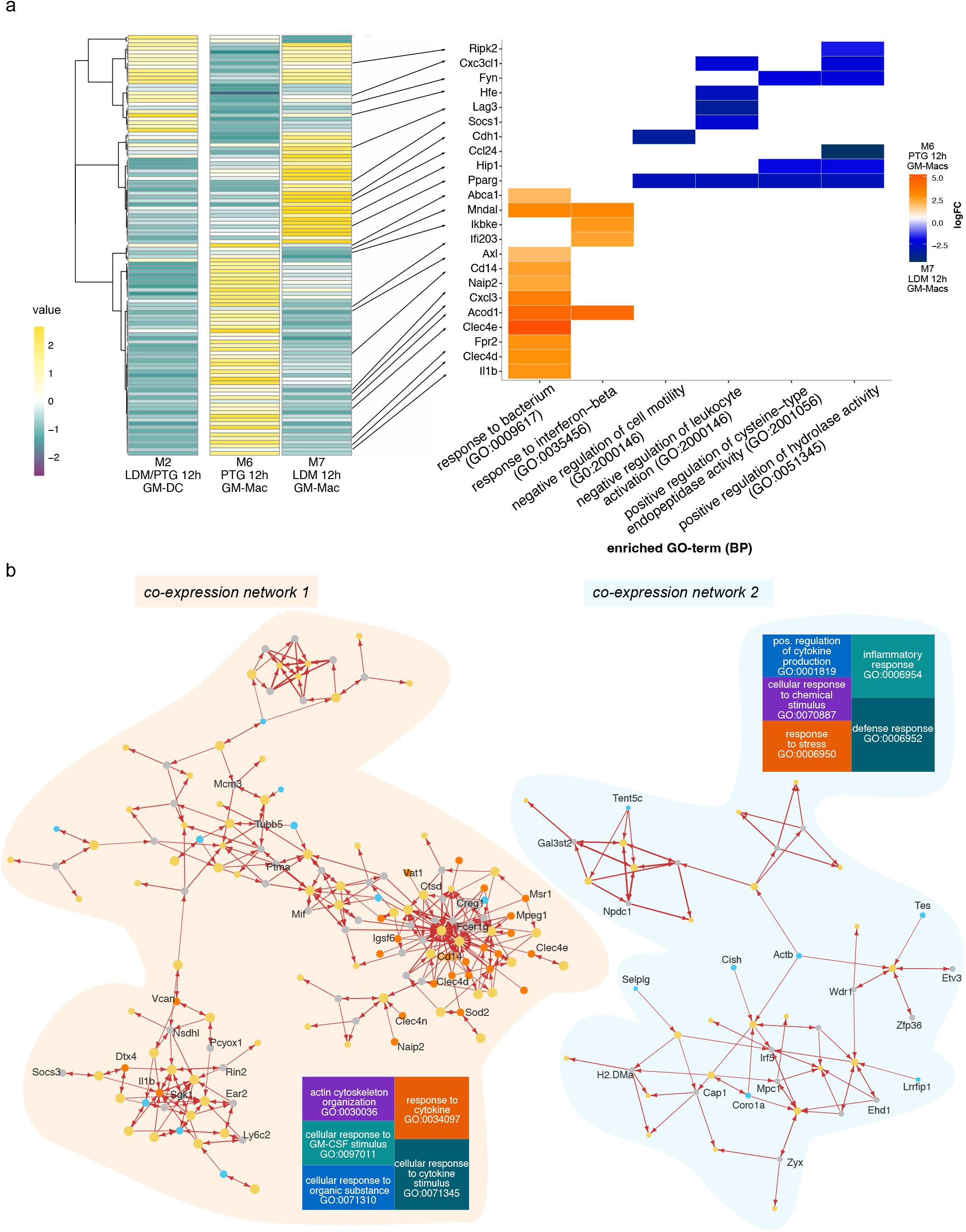
Gene expression and inter-species co-expression patterns of PTG and LDM infected BMDCs 12h post infection. **a)** Heatmap of differentially expressed genes between cluster M6 (mainly GM-Macs containing PTG infected cells 12 hpi) cluster M7 (mainly GM-Macs containing LDM infected cells 12 hpi) across all clusters containing 12 hpi cells, including cluster M2 (mainly containing GM-DCs infected with both, LDM and PTG for 12h) (left panel). Average expression values (values) are depicted in a color scale from negative expression (purple, dark) to positive expression (yellow, light). The right panel depicts a heatplot of the logFC of selected genes (indicated with a black arrow) differentially expressed genes between the investigated groups (M6 and M7) and biological processes (GO). Positive values (orange) indicate upregulation in cluster M6 (PTG, 12 hpi). Negative values (blue) indicate upregulation in cluster M7 (LDM, 12 hpi). **b)** Gene co-expression network of differentially expressed genes of cluster M6 (left panel) and M7 (right panel). Nodes represent genes with size representing weight. Node colors indicate gene source, with M7 marker genes in turquoise, M6 genes in orange, *T. gondii* genes in yellow and remaining *M. musculus* genes in gray. Arrows indicate the direction of co-expression. Increased arrow width indicates higher correlation value. Tree plots show the most highly enriched biological processes of the host. The size of each square represents the enrichment of each term, with larger squares exhibiting higher enrichment.

Conversely, genes upregulated in M7 are involved in “negative regulation of cell motility” (GO:2000146), “positive regulation of cysteine-type endopeptidase activity” (GO:2001056), “positive regulation of hydrolase activity” (GO:positive regulation of hydrolase activity) and “negative regulation of leukocyte activation” (GO:2000146) (figure 2a). The majority of them were reported to be important for the limitation of the immune and inflammatory response such as *Lag3, Cdh1, Socs1*, and *Ccl24* mostly involved in the regulation of cytokine release and signaling ^48–51^. In contrast, we also found *Cxc3cl1* and *Hfe*, which are involved in immune system activation ^52,53^. In addition, genes exhibiting balancing effects on pro- and anti-inflammatory signals (*Rip2k, Pparg*) ^54–56^ showed upregulation.

Moreover, clusters consisting of cells infected with *T. gondii* for 12h, irrespective of cell type or clonal lineage, expressed an S-phase cell cycle signature, in line with previous reports ^57,58^ (Supplementary figure 4). This was further validated by expression of the most prominent cell cycle regulators, namely cyclins a-e and their respective kinases, resembling an S-Phase regulatory state in late stage infected cell clusters (Supplementary figure 5). Together with the effects on expression of immune genes this observation highlights the extensive effects of *T. gondii* on gene expression of the host cell.

Co-expression networks identify which genes tend to show coordinated expression. While they don’t provide causality, they enable identification of potential regulatory genes in investigated conditions ^59^. To investigate co-expression networks between *T. gondii* and host genes across single cells, we selected differentially expressed genes between M6 and M7, identified co-expressed genes using a correlation-based approach and visualized them in a network.

The network analysis revealed several co-expression clusters of variable size (Supplementary figure 6). For simplicity, we selected the two largest clusters, containing the highest numbers of M6 upregulated genes (network 1) or M7 upregulated genes (network 2) for further analysis (figure 2b, supplementary data 3). We observed a higher degree of connectivity for the network mainly consisting of M6 marker genes when compared to the network that only contains M7 marker genes, indicating a first differential effect of infection on the gene expression program between cells infected with PTG (M6) and LDM (M7) parasites.

#### Network 1 (M6, PTG)

Overall, we found that host genes present in interaction network 1 are predominantly involved in cytokine response and stimulation (GO:0034097, GO:0071345) and to a lesser extent, in cellular response to organic substances, GM-CSF stimulation and actin cytoskeleton organization (GO:0071310, GO:0097011, GO:0030038) (figure 2b). In addition, we observed co-expression of several marker genes from cluster M6, including *Clec4d, Clec4e, Cd14* and *Mpeg1* with a small set of *T. gondii* genes (figure 2b). Based on the involvement of these host genes on immune related processes we speculate that the expression of a specific, small set of *T. gondii* genes of network 1 may be sufficient to modulate some of these processes.

#### Network 2 (M7, LDM)

For interaction network 2, we found most genes to be involved in defense and inflammatory response (GO:0006952, GO:0006954) or response to stress, cellular response to chemical stimulus and positive regulation of cytokine production (GO:0006950, GO:0070997, GO:0001819). Host genes in network 2 included *Tes, Tent5c, Coro1a, Selp1g* as well as *Cish, H2-Dma* and *Lrrfip1*. These genes are involved in partially opposing functions for inflammatory and immune responses. They are also reported to be involved in processes such as proliferation and tumor suppression ^60,61^. In interaction network 2, we also observed several *T. gondii* genes directing interferon regulatory factor 5 (*Irf5*), suggesting a potential regulatory role of *T. gondii* genes on *Irf5* expression (figure 2b).

### scDual-Seq enables the identification of putative *T. gondii* genes involved in host-pathogen interactions during late-stage acute infection

We continued to investigate parasite genes co-expressed in each network elucidating on coordinated gene expression between the host and parasite. We used an additional correlation based approach to expand the number of co-expressed *T. gondii* genes involved in coordinating gene expression of network 1 and network 2. In brief, we selected *T. gondii* genes showing positive or negative correlation with *T. gondii* genes found in network 1 or network 2. To reduce the potentially masking effects caused by differences in expression of replication genes of the parasite, we excluded genes of the S/M and G1 sub-transcriptome described using published bulk data ^62^.

We observed that the expanded set of co-expressed *T. gondii* genes of network 1 (ETGN 1), characterized by PTG parasite infection, included a larger number of genes exhibiting mostly positive correlation with only a few anti-correlated genes. The expanded set of co-expressed *T. gondii* genes of network 2 (ETGN 2) contained fewer genes and exhibited an overall positive correlation with one another indicating these genes tend to be expressed together. (figure 3a, Supplementary figures 7-8, supplementary data 4-6).

**Figure 3.**
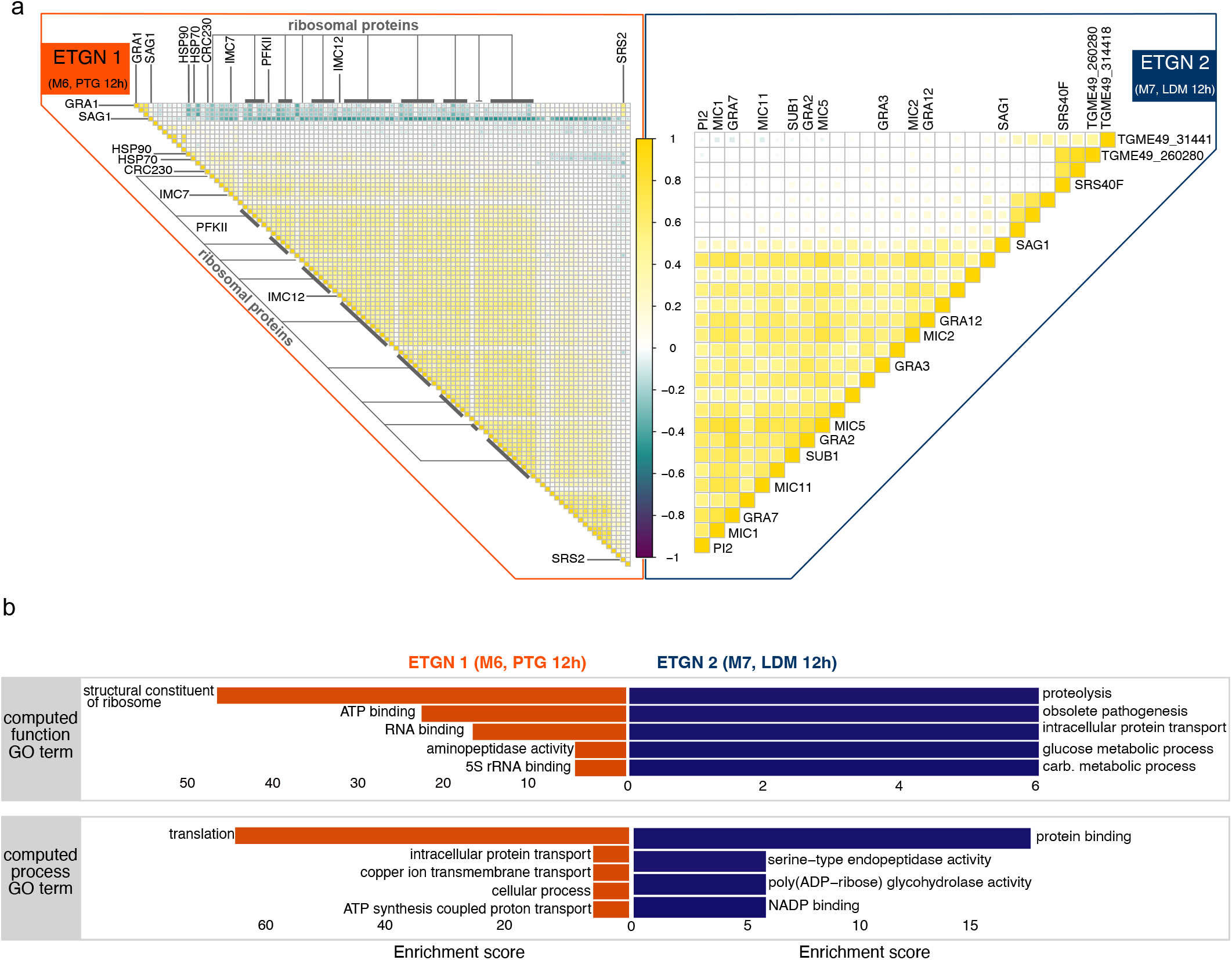
*T. gondii* differential transcription of clonal lineages during infection progression in BMDCs. **a)** Visualization of Pearson correlation coefficients between genes in extended *T. gondii* networks (ETGNs). For *ETGN 1* (left, orange bar) and *ETGN2* (right, blue bar). Correlation values are shown in a gradient from most negative values in dark purple to most positive values in yellow (light). **b)** Gene set enrichment of *T. gondii* genes from correlation analysis shown in **a)** for *ETGN 1* (left, orange) and interaction *ETGN2* (right, blue). The top panel shows enriched functions and the bottom shows enriched processes.

#### ETGN 1 (M6, PTG)

The majority of genes in ETGN 1 (PTG, 12h) belongs to the group of ribosomal proteins. We observed highest correlation values between ribosomal protein genes and genes important for the inner membrane complex (*IMC12* and *IMC7*) the phosphofructokinase *PFKII* as well as a number of genes without annotated function. We also detected positive, but slightly lower correlation values between ribosomal proteins, heat shock proteins, *HSP70* and *HSP90*, as well as *CRC230*, a corepressor complex, suggested to play a role during stage differentiation of the parasite ^63^. We observed a negative correlation between two members of the SRS (SAG1-related sequences) superfamily, namely *SAG1* and *SRS2*, and the majority of the remaining ETGN 1 genes. SRS proteins have been linked to host cell attachment and activation of host immunity to regulate virulence ^64^. Our observations indicate that ribosomal protein genes and SAG-related sequences may fulfill opposite tasks in the parasites (figure 3a, Supplementary figure 7, supplementary data 4-5).

In agreement with the observed correlations of a high number of genes encoding ribosomal proteins, we found high enrichment of structural constituents of ribosome and RNA-/5s rRNA binding function as well as translation processes. Together with the high enrichment of energy metabolism related processes and functions, this may indicate that the PTG parasite’s increased energy expenditure is necessary for survival and the potential preparation for stage differentiation processes such as the formation of bradyzoites (figure 3b).

#### ETGN 2 (M7, LDM)

ETGN 2 (LDM, 12h) is characterized by positive correlation between dense granule (*GRA12, GRA3, GRA2, GRA7*) and microneme *(MIC1, MIC11, MIC5, MIC2, SUB1)* ^40,65^ associated genes. In addition, *PI2*, which was reported to be involved in the control of virulence and differentiation ^66^ shows strong positive correlation between dense granule and microneme associated genes. Conversely to ETGN 1, we found that *SAG1* exhibited a strong positive correlation with the remaining genes of ETGN 2. We found two genes *TGME49_314418* and *TGME49_260280*, which exhibited negative correlation to either only *PI2* or a number of microneme and dense granule associated genes (*MIC1, GRA7, MIC11, SUB1, GRA2* and *MIC5*), respectively (Figure 3a, Supplementary figure 8). While *TGME49_260280* has not been associated with a function, *TGME49_314418* has been described to be a member of the clathrin adaptor complex family ^67,68^ (figure 3a, Supplementary figure 8, supplementary data 4+6).

We observed enrichment of the GO terms for metabolic processes and functions, indicating that the LDM strain, which is overrepresented in ETGN 2, is more effectively adapting to the macrophage environment. Although the term “obsolete pathogenesis” remains controversial given its annotated status as obsolete, we found it appropriate to highlight given its enrichment in more virulent LDM parasites. Thus the obsolete pathogenesis term may be related to virulence or host-pathogen interactions (figure 3b).

### *T. gondii* parasites exhibit gene expression changes within a different cellular compartment within GM-DCs and all identified GM-Mac clusters

We sought to provide further detail on how the parasite modulates its gene expression in response to the host investigating exclusively *T. gondii* gene expression. In contrast to the infected host cells, *T. gondii* parasites were actively replicating. In the PCA, variability along PC1 was attributed to differences in cell cycle stages after comparison to the cell cycle sub-transcriptomes ^62^. We visualized these cell cycle stages on the initial clustering and showed that the identified clusters mainly corresponded to the replicative state of the parasites (Supplementary figure 9). Therefore, we assigned cell cycle stages (S/M or G1) across all cells, using previously described sub-transcriptomes and regressed the cell cycle out of our dataset, resulting in 5 clusters (Tg1-5) across infection conditions (figure 4a,b).

**Figure 4.**
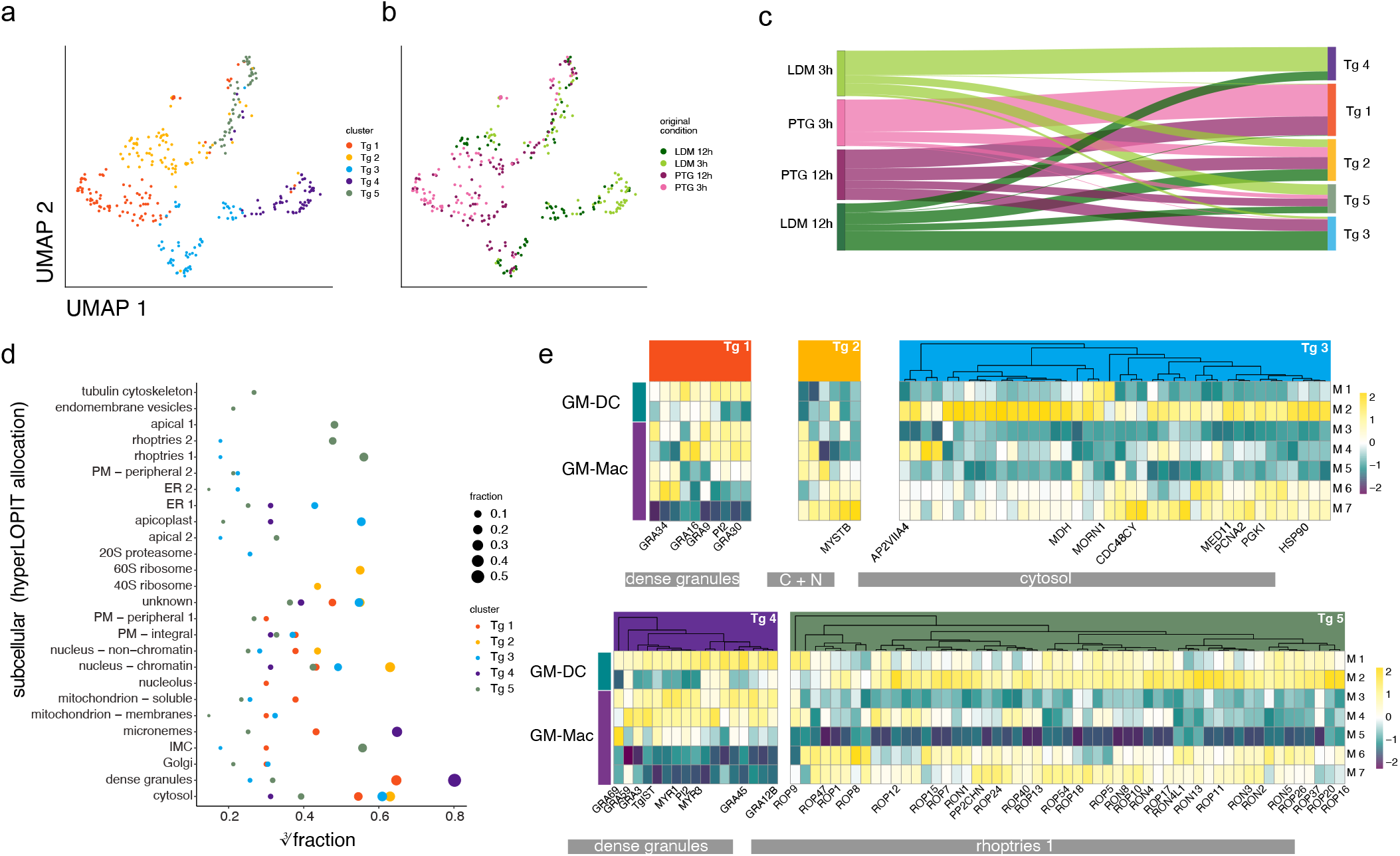
*T. gondii* gene expression profiles and differential association with subcellular localisation across parasite and host clusters. **a)** UMAP projection of *T. gondii* infected cells grouped by identified cluster. **b)** UMAP projection of *T. gondii* infected cells grouped by condition. **c)** Proportion of cells of each original infection condition (left) associated with each identified cluster (right). The color of the ribbon refers to the original condition. The color of the right bar indicates the cluster association (Tg1-Tg5). **d)** Cube root transformed Fraction of marker genes of identified *T. gondii* clusters encoding proteins located to different localisations in *T. gondii* identified previously using hyperLOPIT ^69^. The size of each dot refers to the non-transformed fraction of differentially expressed genes to each location on the y-axis. **e)** Heatmap depicting differentially expressed *T. gondii* genes grouped by *T. gondii* clusters (Tg1-Tg5) across host clusters exhibiting infection (M1-M7). For each *T. gondii* cluster, genes of the highest fraction of intracellular localization were selected for comparison. *T. gondii* clusters are represented in the colors selected for each cluster. Cell type annotations of host clusters are indicated by turquoise (GM-DC) and purple (GM-Mac). Averaged gene expression is shown in a color gradient from low expression (dark purple) to high expression (yellow).

When comparing the infection conditions with the identified clusters, we show that cluster Tg 1 is characterized by PTG specific expression. Cluster Tg 2, similar to Tg 5, is characterized by the presence of parasites of both strains and timepoints. Tg 3 mainly consisted of parasites from 12 hpi, while Tg 4 consisted mainly of LDM parasites (figure 4c).

To further define what is driving the differences between parasites within each cluster, we investigated whether they express organelle-specific genes by comparing our gene expression results to spatially resolved proteome data on the subcellular level ^69^ (figure 4d). While transcription data presented here does not equal protein expression and subsequent effector activity, comparison with spatially resolved proteomics data may give an indication about what processes the parasites are preparing for, providing a glimpse into the future of the parasite’s development.

We found that Tg1 and Tg4, representing the most strain specific clusters for PTG and LDM, respectively, showed the highest fraction of genes belonging to dense granules. The second largest fraction for Tg4 consisted of microneme located genes while the second largest fractions for Tg1 contained genes associated with the cytosol or unknown location. Across clusters containing mixed parasite populations, Tg2, which consists of the highest amount of early stage parasites, exhibits the highest fraction of genes located to the nucleus, the cytosol and the 60S ribosome as well as unknown localization. Tg3, containing mostly late stage parasites, shows similar distributions of genes across localizations as Tg 1, with higher fractions within the apicoplast and the ER (figure 4d). Tg 5 generally shows the highest number of upregulated genes across all *T. gondii* clusters, (Supplementary figure 10, supplementary data 7) with the majority associated with rhoptries, the inner membrane complex (IMC) and the apical region of the parasite (figure 4d, supplementary data 8). Rhoptry proteins, located in the apical region of the parasite, fulfill multiple roles for virulence, host cell invasion and potentially parasitophorous vacuole formation as well as manipulation of host response ^70–72^. The observed differences between these clusters indicate that parasites within each cluster undergo or prepare themselves for different processes during infection, for example invasion, egress, stage differentiation or immune evasion and virulence.

Next we examined the relationship between parasite clusters (Tg 1-5) and their gene expression profiles and *T. gondii* infected host cell clusters (M1-7). For each *T. gondii* cluster (Tg 1-5), we selected *T. gondii* marker genes of the most prominent localization. We observed obvious differences in *T. gondii* expression profiles for markers across host clusters (figure 4e, Supplementary figure 11-12). For instance, when comparing dense granule genes between Tg 1 (PTG) and Tg 4 (LDM) upregulated genes, we observed strong downregulation of all Tg 1 dense granule marker genes in cluster M7 (LDM 12 hpi), while they are partially upregulated (*TGME49_248140, TGME49_266050* and *GRA34*) in M6 (PTG 12 hpi). However, dense granule markers of Tg 4 show general upregulation in GM-Macs and GM-DCs in early time point clusters (M1, M3-M5), with lowest expression in cluster M5. Simultaneously, most dense granule genes are downregulated in all late time point clusters (M2, M6-M7) (figure 4e). In cluster M2 (GM-DCs), we observed the strongest upregulation of rhoptry genes. In addition to observed differences between known subtypes of host cell types (GM-DCs and GM-Macs), we observed downregulation of all rhoptry markers and other location markers of Tg 5 in host cell cluster M5 (figure 4e). For expression of cytosolic and nuclear genes, which describe the highest fraction of upregulated genes in cluster Tg 2 and Tg 3, we observe the strongest differences between time points of infection (M2, M6 and M7 compared to M1 and M3-M5).

### Infection with *T. gondii* shares host response similarities to stimulation by CpG-containing oligodeoxynucleotides (cpGB) and other immune stimulating molecules

Our findings suggest that *T. gondii* LDM (LDM) and PTG (PTG) provoke differential host cell responses primarily in GM-Macs and that an extended network of *T. gondii* specific genes are involved in these host-pathogen interactions at the single cell level. Beyond this, we wanted to explore which innate immune responses could be involved in the differential reactions of the two clonal lineages, comparing host responses to other common microorganisms.

We utilized RNA-seq data from a recent publication by Pandey and colleagues in which they studied the immune response towards seven common microbial signals in BMDCs ^73^. We intersected gene markers of our original experimental conditions and differentially expressed genes for each microbial stimulus in the reference. We observed the highest overlap in upregulated genes stimulated with CpG DNA type B (cpGB) and depleted zymosan, a β-glycan found in fungal cell walls, in both clonal lineages (figure 5a). cpGB is known to target pattern recognition receptors (PRRs) TLR9 ^74^ and Dectin-1 ^75^. Further, we found cells infected with PTG parasites to exhibit higher overlap of expressed genes across most other microbial stimuli (figure 5a). We found *Igsf8* as the only shared marker gene between LPS stimulation and PTG parasite infection, leading us to examine gene expression markers for LPS stimulation in our dataset in more detail. Indeed, we find *Igsf8* expression in both: cells infected with LDM and PTG parasites. However, expression in GM-DCs is only observed for LDM infection. Overall we observed clear differences in expression based on cell type (GM-DC or GM-Mac), and higher expression for some genes in GM-Macs infected with PTG parasites (figure 5b).

**Figure 5.**
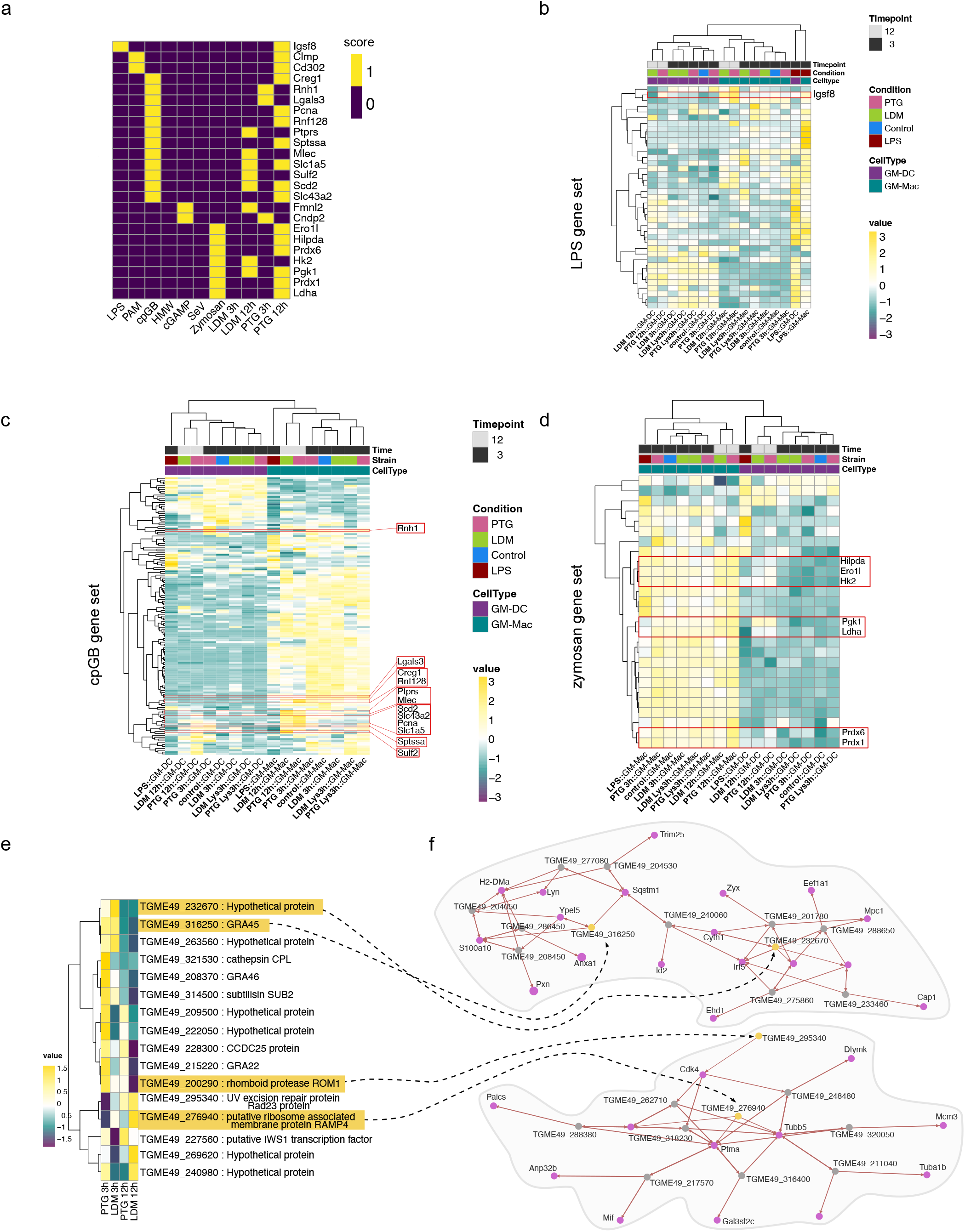
Cell type differentiation and comparative analysis of response to *T. gondii* infection with established PAMPs and *T. gondii* INF_γ_ challenge. **a)** Heatmap depicting commonly expressed genes between BMDCs exposed to established PAMPs (LPS, PAM, cpGB, HMW, cGAMP,SeV and Zymosan) and cells infected with *T. gondii* LDM and PTG for 3 or 12 hours. A score of 1 (yellow) indicates differential expression of the gene in the respective condition while a score of 0 indicates no differential expression. **b)** Heatmap depicting expression of genes upregulated in LPS treated BMDCs ^73^, hierarchically clustered by condition in both cell types and gene expression. Expression values are shown in a gradient stretching from negative expression (dark purple) to positive expression (yellow). The red box highlights expressions of *Igsf8*. **c)** Heatmap exhibiting gene expression values for previously identified cpGB signature and **d)** zymosan signature. Shared marker genes between the reference signature ^73^ and our data are highlighted in red boxes. **e)** Heatmap visualizing conducted comparative analysis between genes identified to be essential during INFγ stimulation in BMDMs ^81^ and their expression profiles in *T. gondii* infective states in our study (LDM and PTG infection 3 and 12 hpi). **f)** Co-expression networks for essential *T. gondii* genes. Genes exhibiting correlation are connected with a black dashed arrow and highlighted in yellow. Correlating host genes are depicted as purple nodes. Remaining *T. gondii* genes are shown as gray nodes. The arrow indicates the direction of the correlation. Increased arrow width indicates higher correlation value.

Based on high similarities of marker expression between *T. gondii* infected conditions and zymosan as well as cpGB stimulated BMDCs, we examined gene expression of these conditions. For cpGB-stimulated BMDCs, we observed higher expression of genes in GM-Macs in comparison to GM-DCs, irrespective of infection. Of those genes that were shared between cpGB challenge and *T. gondii* infection, upregulation was more prominent in PTG infected cells (figure 5c).

Gene expression of marker genes for zymosan challenge was also higher in GM-Macs. Some redox genes (*Prdx1* and *Prdx6)*, genes involved in lipid and cholesterol metabolism (*Hilpda, Er1l)* ^76,77^, stress response and cell death (*Hk2, Pgk1*) ^78,79^ as well as M2 macrophage activation (*Ldha*) ^80^ demonstrated overlap with transcriptional response to zymosan, and showed higher upregulation in both cell types infected with PTG parasites 12 hpi. (figure 5c).

We then sought to use a similar approach to compare *T. gondii* expression in our cell population to other relevant cell population data. Therefore, we employed publicly available data from in vitro CRISPR-Cas9 screens of *T. gondii* genes comparing it to our dual-scSeq data. When we intersected *T. gondii* marker genes of our infected conditions with high confidence hits for “fitness under INFγ stress” identified by Wang et al. ^81^, we found the majority of intersecting genes to be upregulated in parasites 3 hpi, with PTG strains upregulating a higher number of genes compared to LDM strains. We also found genes TGME49_276940 and TGME49_269620, exclusively upregulated in LDM parasites 3 hpi or 12 hpi, respectively (figure 5d).

To further characterize interactions between *T. gondii* INFγ stress marker genes and host genes, we generated co-expression networks, resulting in multiple networks of variable sizes (Supplementary figure 13). For the two largest networks, we found *GRA45* (*TGME49_316250*), upregulated in PTG parasites at 3 hpi, to be co-expressed with *YpeI5, S100a10* and *Sqstm1*, which are all reported to be involved in regulation of innate immune response and inflammation ^82,83^ as well as autophagy in the case of *Sqstm1* ^84^ (figure 5e). Conversely, *TGME49_276940* and *TGME49_295340*, upregulated in LDM parasites, showed co-expression with *Tubb5, Ptma* or *Cdk4*, respectively (figure 5f). *Tubb5* and *Ptma* are suggested to play a role in filamentation and pathogenesis of *Streptococcus agalactiae in vitro* ^85^ or *Candida albicans* ^86^, respectively.

Moreover, co-expression of *Tubb5* and *Cdk4* with essential *T. gondii* genes corroborates the suggested manipulation of the cell cycle by *T. gondii*.

## DISCUSSION

*T. gondii* is a successful intracellular parasite establishing interacting with a variety of host cells including; macrophages and dendritic cells. In this study, we performed scDual-Seq of BMDCs infected with two phenotypically distinct clonal lineages of *Toxoplasma gondii*. Our results demonstrate the importance and value of scDual-Seq approaches to investigate host-pathogen interactions of heterogeneous populations of infected cells and provide a resource for future studies on host-pathogen interactions.

We analyzed infection dynamics over two timepoints (3 and 12 hpi) during acute *T. gondii* infection. Dual-scSeq enabled us to describe expression patterns of individual populations of host cells in tandem with *T. gondii* gene expression, parsing out heterogeneity in host-pathogen interactions and potential genes involved in different host-pathogen interaction programs over time. Information on heterogeneity of host-pathogen interactions was lost in previous bulk-RNA sequencing studies on *T. gondii* infected BMDCs. By using a scSeq approach we were able to show that the parasite manipulates two subpopulations of infected BMDCs, GM-Macs and GM-DCs differently. ^25–27^.

Our data indicates that only the GM-Mac subpopulation 12 hpi showed differential expression profiles based on the infecting parasite strain. This suggests that this population of cells is either more susceptible to strain specific modulation by PTG and LDM parasites or that the GM-Mac subpopulation specifically responds to different parasite strains. This highlights the importance of considering transcriptional heterogeneity of host-pathogen interactions, previously suggested by single cell studies on human monocytes infected with *T. gondii* parasites ^31^. However, this study investigates infection dynamics at 1 hpi and does not investigate transcriptional differences between potential subpopulations of host cells infected with parasite strains of different genetic background or the corresponding parasite expression dynamics. Thus, our data significantly augments existing data, including multiple time points of infection, different subpopulations of murine host immune cells as well as responses to the two *T. gondii* genotypes with highest global prevalence.

Previous studies describe transcriptional differences between GM-DCs and GM-Macs at steady-state and after exposure to LPS ^5^. Our study expands on the observed differences by providing single cell resolution in our dataset, including our control population of LPS treated BMDCs.

However, the main point of interest in our study includes the transcriptional differences induced in the BMDCs during acute *T. gondii* infection. Upregulation of genes associated with anti-inflammatory effects in LDM parasite infected GM-Macs as opposed to upregulation of a pro-inflammatory response in PTG infected cells is in agreement with previous studies in murine macrophage infections ^25^. The higher similarity of the gene expression signature of anti-inflammatory signals in GM-DCs infected with *T. gondii* parasites of the LDM as well as the PTG strain, might suggest that the parasite exploits or even promotes immune tolerogenic capacities of infected DCs. DCs represent important regulators of the balance between tolerance and immunity, giving rise to large numbers of subtypes partially exhibiting immune tolerogenic phenotypes ^87,88^. In addition, stimulation with GM-CSF in BMDCs was shown to favor immune tolerogenic DC populations ^89,90^. This suggests that in addition to the reported manipulation of the migratory behavior of DCs, promoting parasite dissemination ^16^ the parasite may be more efficient in generating a suitable environment for replication and survival in GM-DCs in comparison to GM-Macs. Conclusively, our observations highlight the importance of considering heterogeneity in host-pathogen interactions and prompt further, more detailed studies on interactions between *T. gondii* and the described subsets, particularly of GM-DCs.

To investigate differential host-pathogen interactions between parasites and GM-Macs after 12 hpi in more detail and understand the reason for differential responses to infecting parasites, we generated and examined co-expression networks of host- and parasite gene expression based on differential marker genes of cluster M6 (PTG, 12h) and M7 (LDM, 12h). We interpret the higher connectivity in *network 1* to be indicative of an increased response to parasite expression in PTG parasites at 12 hpi in infected GM-Macs. Genes exhibiting strong correlation with only a few parasite genes within the network included C-type lectins *Clec4d, Clec4e* and *Clec4n*, important players in innate recognition of pathogens and interleukin production as well as phagocytosis of parasites in macrophages ^43,91,92^. Further, *Cd14*, glycosylphosphatidylinositol (GPI)-anchored receptor, which serves as a co-receptor for several Toll-like Receptors (TLRs) ^44^, showed co-expression with a tight cluster of genes expressed by PTG parasites and high correlation with the expression of *T. gondii* SRS2 (TGME49_233480). SRS2 has been implicated with IL-10 release and thus anti-inflammatory activity, interacting with MIC1 and MIC4 ^44^. The co-expression of these genes may be crucial for the formation of bradyzoites in PTG parasites ^93^ as opposed to LDM parasites, which are less likely to form mature bradyzoites ^94^.

Genes within network 2 involved in immune system processes included *Coro1a* and *Selp1g* associated with leukocyte trafficking during inflammation ^95,96^ and implications in host cell modulation of macrophages infected with *Staphylococcus aureus* ^97^ or the restriction of effector T cell responses in acute viral infection ^96^. This supports the hypothesis that *T. gondii* LDM parasites modulate host cell responses and potentially limit them more effectively than PTG parasites. Apart from immune system related genes we also find a number of genes involved in tumor suppression including *Tes* ^60^ and *Tent5c* ^61^. Together with ours and previous observations ^57,58^, that *T. gondii* changes the cell cycle expression profile toward an S-phase state upon infection progression, indicate that *T. gondii* infection particularly with lineages of RH descent, might have a more global effect on host cell proliferation and cell cycle dysregulation than reported so far.

The observed positive correlation between many genes encoding ribosomal proteins and a large number of host genes involved in energy expenditure, phagocytosis and inflammation in extended *T. gondii* network 1 (*ETGN 1*) suggests that the majority of parasites may express these genes in response to the pro-inflammatory immune response of the infected host cells. Moreover, the highly expressed ribosomal proteins may represent an interesting pool of potent targets for the innate immune system. For instance, ribosomal protein P2 has been previously identified as a target for protection against toxoplasmosis ^98^.Based on previous reports of the importance of MIC1, MIC2 and GRA2 for in effective immune evasion or virulence ^40,99,100^, *T. gondii* genes encoding secreted proteins in *ETGN 2* likely belong to a network of genes with importance for these processes. Conclusively, while not providing causality or evidence for direct interactions, which is beyond the scope of this study, we consider our findings a valuable resource to investigate potential direct interactions between genes that correlate positively in expression in single cells for future functional studies on host-pathogen interactions.

Early stage parasites of both lineages showed expression of dense granules and micronemes, known to be important during cell adhesion and invasion of the parasite ^101^. Recent studies show that dense granules can alter the behavior of host cells directly, as shown for altered cell-motility by GRA28 in infected macrophages ^17^.

The overall high fraction of dense granule genes at 3 hpi and the differential expression of these genes between LDM and PTG parasites further support the importance of dense granules for host-pathogen interactions and potential strain-specific host-cell modulation during the subsequent parasite development. Comparing the expression of location specific genes across host-cell clusters, we observed the absence of all genes encoding for rhoptry proteins in one sub-cluster of GM-Macs. Multiple rhoptries have been implicated in virulence and modification of host response, such as ROP16 and ROP18 ^72,102^, indicating that parasites in this sub-cluster are modified differently or *T. gondii* parasites are less effective in establishing virulence, potentially being successfully targeted by the immune system. It is important to note that transcription data presented here does not equal protein expression and subsequent effector activity. Thus our data draws an image of the potential future development of the parasite within a host-cell.

Comparing gene expression of *T. gondii* infected host cells to expression profiles of cells stimulated with a variety of PAMPs, we found the highest expression similarities to cpGB as well as zymosan stimulation, primarily in PTG infected GM-Macs 12 hpi. Zymosan is a ligand for TLR2 and Dectin-1 in macrophages and dendritic cells. Interactions between zymosan and host PRRs eventually leads to the secretion of pro-inflammatory cytokines (TNF-alpha, IL-8) and activation of the Dectin-1/Syk/NF-□B signaling pathway which mediates an inflammatory response ^103^. CpG motifs have been mainly implicated with B-cell function ^74,104,105^ and are described to be ligands for TLR9, leading to the activation of the NF-LB-REl dependent innate pathway, cooperating with IL-10, STAT proteins and IFN-responsive factors ^105^. *T. gondii* infection stimulates INF production and pro-inflammatory cytokine production in a Myeloid Differentiation factor 88 (MyD88) dependent manner. It is proposed that this pathway is mainly activated through stimulation of TLR11 ^39,106^. However, there is additional evidence of the involvement of TLR9 ^106,107^ and TLR2 ^106,108^, confounded by similarities in expression patterns in our data. Moreover our data provides gene lists of the most important genes expressed during a potential TLR2 or TLR9 stimulation by *T. gondii* parasites of different clonal lineages.

In line with our observations, the PTG lineage shows stronger activation of TLR2 and TLR9 related gene expression profiles we find the highest overlap and upregulation of essential *T. gondii* IFNγ response genes during early timepoints (3 hpi) and in parasites of the PTG lineage. Co-expression analysis revealed genes involved in modulation of innate immune responses and TLR function are co-expressed with dense granule gene, GRA45, implying a potential role in immune modulation during an inflammatory response of the host. In addition, we find co-expression of genes related to cytoskeleton formation and pathogenesis as well as cell cycle regulation, supporting the diversity of host cell modulation by *T. gondii*.

In summary, this study provides a detailed resource of host-pathogen interactions over the time-course of an acute *T. gondii* infection in BMDCs. The identification of differential responses in distinct sub-populations of cells highlights the importance of studying infection kinetics at the resolution of single infected cells and raises the question how and why parasites of different clonal origin modulate cell types differently or change cell type signatures. We show co-expression of parasite ribosomal proteins and pro-inflammatory genes in PTG infected cells and of dense granules and immune modulating genes in LDM infected cells, highlighting differential modulation of host-pathogen interaction pathways between clonal lineages for the first time at the single-cell level. Further, we show differential expression of rhoptry proteins, dense granules and micronemes between PTG and LDM parasites as well as clusters of host cells, with strain specific host modulation by dense granules and micronemes while rhoptry expression is similar between strains. This highlights the importance of dense granules for nuanced host-pathogen interactions, prompting future functional studies on these effectors. Moreover, the integration and comparative analyses between our data and previously published data highlights the possibility of validating and identifying new expression programs during host-parasite interactions.

In conclusion, we anticipate that the data presented here provides a unique and comprehensive resource for studies on host-parasite interactions between *T. gondii* parasites and infected immune cells. We further anticipate that the resource that we provide herein serves as a tool which can be applied to study host-pathogen interactions on a variety of other invasive pathogen and host cell complexes of medical and veterinary importance, including *Plasmodium, Cryptosporidium, Babesia*, and *Leishmania* as well as different bacteria and viruses.

## METHODS

### Cell culture

Primary bone marrow-derived mononuclear phagocytes were generated as previously described ^109,110^. Briefly, cells from the bone marrow of 6-10 week old C57BL/6 mice (Charles River) were cultured in RPMI 1640 (Gibco) complemented with 10% fetal bovine serum (FBS, Sigma), gentamicin (20 μg/ml; Gibco), glutamine (2 mM; Gibco) and HEPES (0.01 M; Gibco), referred to as complete media, and further supplemented with 10 ng/ml of recombinant mouse granulocyte-macrophage colony stimulating factor (GM-CSF) (Peprotech). Culture media was replenished on days two, four and six. Loosely adherent cells^111^ were harvested on day seven or eight for experiments.

The parasite lines used include GFP-expressing RH-LDMluc (LDM, cloned from RH-GFPS65T) and GFP-expressing PTGluc (PTG, cloned from ME49/PTG-GFPS65T) ^112^. Tachyzoites were maintained by serial 2-day passaging in human foreskin fibroblast (HFF-1 SCRC-1041, American Type Culture Collection) monolayers cultured in Dulbecco’s modified Eagle’s medium (DMEM; Gibco) with 10% FBS, gentamicin (20μg/ml), glutamine (2mM), and HEPES (0.01 M).

### Infection challenges, lysates and LPS

On day seven or eight, BMDCs in complete media were challenged with freshly egressed *T. gondii* tachyzoites. To ensure an infection-rate of single parasites above 20%, cells at 3 hours were infected at a multiplicity of infection (MOI) 3, or 12 hours at MOI 1.5. Negative controls were treated with parasite lysates or not stimulated, while positive controls were treated with 100ng/ml LPS for 3 hours. The parasite lysates were produced by ultrasonication of live tachyzoites, where ultrasonication was applied 3 times at 10 second intervals with 1-minute interval on ice.

Post-infection, cells were treated with Fc-block (anti-mouse CD16/CD32 antibody, 1:200; BD 553142) for 15 minutes, washed with D-PBS and further incubated with a monoclonal antibody against CD11c conjugated with PE-Cyanine7 (1:200; Thermofisher 25-0114-82) for 30 minutes, washed with D-PBS and resuspended in ice-cold D-PBS. Cells were kept on ice prior to sorting.

### Cell sorting and library preparation

A preliminary trial was set up to validate optimal cycling conditions for cDNA amplification of single *T. gondii-*infected BMDCs by performing rt-qPCR of target genes known to be expressed in *T. gondii* and genes known to be expressed in DCs (primer sequences).

The sorting was performed with a MoFlo Astrios EQ (Beckman Coulter, USA) cell sorter using 488 and 532 nm lasers for excitation, 100 μm nozzle, sheath pressure of 25 psi and 0.1 μm sterile-filtered 1 x PBS as sheath fluid. Flow sorting data was interpreted and displayed using the associated software, Summit v 6.3.1.

To test the precision of the adjustments made to center the drop in each well, a colorimetric test mimicking the sort was done based on ^113^. A 1.5 mg/μl solution of HRP (cat no 31490, ThermoFisher Scientific) with 1 drop of flow check beads (Beckman Coulter, USA) was sorted into each well of an Eppendorf 384-well plate (Cat no 34028, ThermoFisher Scientific). A color change after sorting indicated that the drop hit the sort buffer and that the precision was adequate.

Individual BMDCs or single *T. gondii* infected BMDCs were deposited into 384-well plates (Eppendorf twin.tecTM PCR plates) containing 2.3 μl of lysis buffer ^114^ using a CyClone™ robotic arm and at highly stringent single cell sort settings (single mode, 0.5 drop envelope). Side scatter was used as the trigger channel and sort regions were based on *Toxoplasma* cells expressing GFP and the surface antibody CD11C-PE-Cy7 bound to human dendritic cells. GFP was excited at 488 nm and detected with a 530/40 nm bandpass filter whereas PE-Cy7 was excited at 532 nm and detected using a 695/70 nm bandpass filter. The plate and sample holder were kept at 4 ºC at all times during the sort. After the sort, the plates were immediately spun down and put on dry ice.

Single BMDCs or single *T. gondii* infected BMDCs were sorted directly into lysis buffer and cDNA libraries were generated using a slightly modified version of Smart-seq2 as previously described ^114^, but where we used 20 cycles for cDNA amplification.

### Single cell sequencing

Single-cell libraries were sequenced at the National Genomics Infrastructure, SciLifeLab Stockholm, using the HiSeq2500 platform (Illumina) for 56 bps single-end sequencing. We sequenced a total of 764 (negative controls n=2, per plate) single BMDCs or *T. gondii* infected BMDCs from a total of 8 different conditions.

## Computational analysis

### Mapping and annotation and filtering

A custom reference genome was made by combining *Mus musculus* GRCm38 and *Toxoplasma gondii* TGA4.44. The reads were aligned to the genome using STAR v 2.7.2, and gene expression was measured using featureCounts v 2.0.0, using default settings. Cells with less than 10 000 mapped reads were filtered due to substantially inferior quality and the remaining 518 cells were subjected to subsequent computational analysis. A total of 62 276 genes across 532 cells for the host (*Mus musculus)* and *T. gondii* were analyzed.

### Normalization, dimensionality reduction and clustering

Main computational analysis of read-count matrices was performed using the Seurat package (v 4.0.3) ^115^ in R (v 4.2.0). The complete R workflow can be assessed in an R markdown (see code availability section). First, count matrices and metadata were loaded and split by the respective species. Ensembl IDs of genes were translated to gene symbols and cells with a mitochondrial gene count above 10% were filtered. Subsequently reads were normalized for sequencing depth using the “SCTransform” function in Seurat, selecting the top 3000 variable genes ^116^.

Thereafter, dimensionality reduction was performed using PCA, computing the first 50 PCs. The first 15 PCs from the analysis were then subjected to shared-nearest-neighbor (SNN) inspired graph-based clustering via the “FindNeighbors” and “FindClusters” functions. For modularity optimization, the louvain algorithm was used and clustering was performed at a resolution of 0.8 for clustering granularity, resulting in 9 clusters. After clustering, a UMAP dimensionality reduction was performed.

### Differential gene expression analysis (DGEA)

Differential gene expression analysis (DGEA) of genes in identified clusters was performed using the function “FindAllMarkers” from the Seurat package (v. 4.0.5). Following the default option of the method, differentially expressed genes for each cluster were identified using a non-parametric Wilcoxon rank sum test. Differentially expressed genes in a cluster were defined by setting initial thresholds above a logarithmic fold-change of 0.5 and being present in at least 25% of the cells belonging to the same cluster. Representative marker genes with an adjusted *p*-value below 0.05 for each cluster were further selected. *p*-values were adjusted using a Bonferroni correction including all genes in the dataset. To find representative marker genes with elevated expression in comparison to the remaining clusters, only positive log fold-changes were considered. For individual analyses such as gene enrichment analysis (see “Gene set enrichment analysis (gsea)”), threshold values for differential gene expressions were modified and will be described in detail in the respective sections of the materials and methods and results. To identify DEGs between specific clusters of interest, the “FindMarkers” function in Seurat was used and the identities were set to the respective clusters of interest. The same thresholds as stated above were used to define DEGs.

For visualization purposes of DGE data, the gene expression data for all cells was averaged and grouped according to their cluster identity, resulting in average expression of each gene and cluster. Then, expression data was scaled and clipped to average expression values between −1.5 and 1.5, with negative values representing downregulation and positive values representing upregulation of each gene.

### Cell type classification and annotation

We classified cell types in our data using the clustifyr package (v.1.8.0) ^117^ and the sorted microarray expression data presented in ^5^ as reference. In brief, clustifyr adopts correlation-based methods to find reference transcriptomes with the highest similarity to query cluster expression profiles. After converting the microarray data in the correct format for automated cell type annotation, we used the default settings (Spearman rank correlation) to estimate correlation coefficients between the single cells in our clusters and the reference cell types.

### Cell cycle scoring

To chart the cell cycle phases of individual cells, cell cycle scoring was performed using the “CellCycleScoring” function in Seurat. Scoring was performed based on the host (*M. musculus*) expression data as well as on the *T. gondii* data. Here the scoring function was performed using the default parameters. In brief, the function assigns each cell a score based on its expression of G2/M and S phase markers, which are assumed to be anticorrelated in their expression levels. Thus, cells expressing neither are assumed to be in G1 phase. Cell cycle associated marker gene lists for mouse were retrieved from: https://raw.githubusercontent.com/hbc/tinyatlas/master/cell_cycle/Mus_musculus.csv (20210707). These gene lists are based on orthology analyses of human cell cycle genes presented in ^118^.

*T. gondii* cell cycle associated genes were extracted from supplementary data of ^62^. As the cell cycle stages of *T. gondii* contain a S/M and G1 phase only, the resulting phases were changed accordingly.

### Gene set enrichment analysis (GSEA)

To interpret gene expression data and differentially expressed genes further, we performed a gene set enrichment analysis for clusters of interest. This was done using the gseGO function of the clusterProfiler package in R (v.4.0.5). To identify significantly enriched genes, we set the ontology parameter to use gene set enrichment for biological processes and set the gene set parameters to include a minimum of three genes and a maximum of 800 genes in one set. The reference dataset for *M*.*musculus* was used, and the resulting p-value from the fast preranked gene set enrichment analysis (fgsea) was adjusted using a Benjamini-Hochberg correction. For differentially expressed genes between cluster M6 and cluster M7, correction of gene set enrichment resulted in non-significant p-values, which is why we don’t report significant differences for gene set enrichment but only fold changes of genes of interest with their respective biological process (figure 2a).

To identify *T. gondii* specific gene set enrichment we used the ToxoDB database to annotate all *T. gondii* genes present in our dataset and included computed GO terms for I) function and II) process generated by VeuPathDB utilizing InterPro-to-GO ^119^. Then the counts for each computed GO-term within the gene list of interest were determined. After the fraction of genes of the geneset of interest was determined and divided by the fraction of the reference (all data) to determine the enrichment for each term and geneset. We termed the resulting value “enrichment score” in our data.

### Comparative analysis of gene expression and reference data

For reference data integration and comparison we included three publicly available datasets, to I) perform cell type annotations (cell type classification and annotation) ^5^ II) to compare the response in our data to data investigating PRR agonists ^73^ and III) to compare our results to identified essential genes for *T. gondii* during INFγ stress ^81^. For II) we first identified differentially expressed genes for each single condition (PAMP) using DESeq2 v.1.36.0 ^120^. We investigated expression patterns for II) by intersecting genes identified to be upregulated in BMDCs during stimulation with each agonist. For simplicity we generated bimodal scores resulting in two modes: expression (1) and no expression (0). We further show gene expression of PRR agonists by investigating aggregated expression of individual genes across our experimental conditions, visualized in a heatmap. We performed similar analyses for III), additionally investigating co-expression of shared genes between the reference data and our gene expression data.

### Co-expression network analysis

To investigate inter-species co-expression networks we first performed an asymmetric Biweight midcorrelation (bicor) to identify gene expression correlations between *T. gondii* and *M. musculus* genes across cells. Using a k-nearest neighbor approach, we defined the top 20 interactors for each gene in the *T. gondii* expression matrix and *M. musculus* expression matrix. Using igraph (v.1.3.1), we then constructed correlation network plots (Co-expression networks). Using the three nearest neighbors (interactors), a correlation threshold of 0.3/-0.3 and 3 steps. The number of steps defines how many additional interactors after the primary interaction should be displayed after the interactors of the provided seed genes. We displayed the largest interaction clusters using the “induced_subgraph” function in igraph. To integrate publicly available data on *T. gondii*, we investigated co-expression of *T*.*gondii* genes, which were previously shown to be essential during INFγ stress in BMDMs ^81^.

### Generation of extended *T. gondii* co-expression

To investigate gene expression correlation between genes resulting from the co-expression network analysis and other genes in the dataset, we first performed an asymmetric bicor correlation analysis between all genes present in the co-expression network cluster of interest and all remaining genes to determine which genes exhibit positive or negative expression correlation across cells. Then, we decided for an appropriate cut-off, only considering genes to be positively or negatively correlated based on the distribution of correlation values, resulting in thresholds between 0.15/-0.15 and 0.25/-0.25. After identifying a subset of genes exhibiting correlation above or below the determined threshold, we calculated pearson correlations between genes of interest and the subset of correlated genes and visualized them using the corrplot package (v.0.92).

## Data availability

The raw sequencing data will be made available in the Sequence Read Archive (SRA) with the accession number PRJNA918538 and interactively on the VeuPathDB database upon publication.

## Code availability

The code used for the analysis will be made available on Github via https://github.com/ANKARKLEVLAB upon publication.

## Author contributions

**JA, JH** and **ABa** conceived the study. **JA** and **JH** supervised the study. **ABh** cultured cells and performed infections. **JA** and **AMD** performed sorting and scSeq. **MM** and **JH** performed mapping and initial computational analyses. **FH** performed computational analysis of the data and generated figures. **FB** and **FH** designed and implemented co-expression network analysis. **FH, AD** and **JA** wrote the manuscript. All authors read and reviewed the manuscript.

## Acknowledgements

We thank the Microbial Single Cell Genomics and the Eukaryotic Single Cell Genomics facilities at Science for Life Laboratory (SciLifeLab), Sweden, for cell sorting as well as smart-seq2 library preparations, respectively, and the National Genomics Infrastructure, SciLifeLab for sequencing. The computations were performed using resources provided by SNIC through Uppsala Multidisciplinary Center for Advanced Computational Science (UPPMAX). We thank Moritz Treeck and the Treeck group for their scientific advice on *Toxoplasma* biology. **JA** is supported through grants from the Swedish Research Council (VR 2021-05057), and the Swedish Foundation for Medical Research (SSMF). **JH** and **MIMS** are supported by Vetenskapsrådet (VR 2021-06602). **ABh** was supported by a grant from the Swedish Research Council (VR 2018-0241 to ABa). **ABa** was funded by VR 2022-00520. **FH** is in part funded by the Sven and Lily Lawski foundation.

## REFERENCES

1. Medzhitov, R. (2007). Recognition of microorganisms and activation of the immune response. Nature 449, 819–826. 10.1038/nature06246.

2. Hoebe, K., Janssen, E., and Beutler, B. (2004). The interface between innate and adaptive immunity. Nat Immunol 5, 971–974. 10.1038/ni1004-971.

3. Guilliams, M., Ginhoux, F., Jakubzick, C., Naik, S.H., Onai, N., Schraml, B.U., Segura, E., Tussiwand, R., and Yona, S. (2014). Dendritic cells, monocytes and macrophages: a unified nomenclature based on ontogeny. Nat Rev Immunol 14, 571–578. 10.1038/nri3712.

4. Hirayama, D., Iida, T., and Nakase, H. (2017). The Phagocytic Function of Macrophage-Enforcing Innate Immunity and Tissue Homeostasis. Int J Mol Sci 19, E92. 10.3390/ijms19010092.

5. Helft, J., Böttcher, J., Chakravarty, P., Zelenay, S., Huotari, J., Schraml, B.U., Goubau, D., and Reis e Sousa, C. (2015). GM-CSF Mouse Bone Marrow Cultures Comprise a Heterogeneous Population of CD11c+MHCII+ Macrophages and Dendritic Cells. Immunity 42, 1197–1211. 10.1016/j.immuni.2015.05.018.

6. Shalek, A.K., Satija, R., Adiconis, X., Gertner, R.S., Gaublomme, J.T., Raychowdhury, R., Schwartz, S., Yosef, N., Malboeuf, C., Lu, D., et al. (2013). Single-cell transcriptomics reveals bimodality in expression and splicing in immune cells. Nature 498, 236–240. 10.1038/nature12172.

7. Smith, T.G., Walliker, D., and Ranford-Cartwright, L.C. (2002). Sexual differentiation and sex determination in the Apicomplexa. Trends in Parasitology 18, 315–323. 10.1016/S1471-4922(02)02292-4.

8. Flegr, J., Prandota, J., Sovičková, M., and Israili, Z.H. (2014). Toxoplasmosis--a global threat. Correlation of latent toxoplasmosis with specific disease burden in a set of 88 countries. PLoS One 9, e90203. 10.1371/journal.pone.0090203.

9. Dubey, J.P. (2010). Toxoplasmosis of animals and humans (CRC Press).

10. Mousavi, P., Mirhendi, H., Mohebali, M., Shojaee, S., Keshavarz Valian, H., Fallahi, S., and Mamishi, S. (2018). Detection of Toxoplasma gondii in Acute and Chronic Phases of Infection in Immunocompromised Patients and Pregnant Women with Real-time PCR Assay Using TaqMan Fluorescent Probe. Iran J Parasitol 13, 373–381.

11. Hu, K., Johnson, J., Florens, L., Fraunholz, M., Suravajjala, S., DiLullo, C., Yates, J., Roos, D.S., and Murray, J.M. (2006). Cytoskeletal Components of an Invasion Machine—The Apical Complex of Toxoplasma gondii. PLoS Pathog 2, e13. 10.1371/journal.ppat.0020013.

12. Kato, K. (2018). How does Toxoplama gondii invade host cells? J Vet Med Sci 80, 1702– 1706. 10.1292/jvms.18-0344.

13. Barragan, A., and Sibley, L.D. (2002). Transepithelial Migration of Toxoplasma gondii Is Linked to Parasite Motility and Virulence. Journal of Experimental Medicine 195, 1625– 1633. 10.1084/jem.20020258.

14. Delgado Betancourt, E., Hamid, B., Fabian, B.T., Klotz, C., Hartmann, S., and Seeber, F. (2019). From Entry to Early Dissemination—Toxoplasma gondii’s Initial Encounter With Its Host. Front. Cell. Infect. Microbiol. 9, 46. 10.3389/fcimb.2019.00046.

15. Bhandage, A.K., Kanatani, S., and Barragan, A. (2019). Toxoplasma-Induced Hypermigration of Primary Cortical Microglia Implicates GABAergic Signaling. Front. Cell. Infect. Microbiol. 9, 73. 10.3389/fcimb.2019.00073.

16. Lambert, H., Hitziger, N., Dellacasa, I., Svensson, M., and Barragan, A. (2006). Induction of dendritic cell migration upon Toxoplasma gondii infection potentiates parasite dissemination. Cell Microbiol 8, 1611–1623. 10.1111/j.1462-5822.2006.00735.x.

17. ten Hoeve, A.L., Braun, L., Rodriguez, M.E., Olivera, G.C., Bougdour, A., Belmudes, L., Couté, Y., Saeij, J.P.J., Hakimi, M.-A., and Barragan, A. (2022). The Toxoplasma effector GRA28 promotes parasite dissemination by inducing dendritic cell-like migratory properties in infected macrophages. Cell Host & Microbe 30, 1570–1588.e7. 10.1016/j.chom.2022.10.001.

18. Ueno, N., Lodoen, M.B., Hickey, G.L., Robey, E.A., and Coombes, J.L. (2015). Toxoplasma gondii Linfected natural killer cells display a hypermotility phenotype in vivo. Immunol Cell Biol 93, 508–513. 10.1038/icb.2014.106.

19. Lambert, H., and Barragan, A. (2010). Modelling parasite dissemination: host cell subversion and immune evasion by Toxoplasma gondii. Cellular Microbiology 12, 292– 300. 10.1111/j.1462-5822.2009.01417.x.

20. Sibley, L.D., and Ajioka, J.W. (2008). Population Structure of Toxoplasma gondiiL: Clonal Expansion Driven by Infrequent Recombination and Selective Sweeps. Annu. Rev. Microbiol. 62, 329–351. 10.1146/annurev.micro.62.081307.162925.

21. Howe, D.K., and Sibley, L.D. (1995). Toxoplasma gondii Comprises Three Clonal Lineages: Correlation of Parasite Genotype with Human Disease. The Journal of Infectious Diseases 172, 1561–1566.

22. Sibley, L.D., and Boothroyd, J.C. (1992). Virulent strains of Toxoplasma gondii comprise a single clonal lineage. Nature 359, 82–85. 10.1038/359082a0.

23. Kanatani, S., Uhlén, P., and Barragan, A. (2015). Infection by Toxoplasma gondii Induces Amoeboid-Like Migration of Dendritic Cells in a Three-Dimensional Collagen Matrix. PLoS One 10, e0139104–e0139104. 10.1371/journal.pone.0139104.

24. Jensen, K.D.C., Wang, Y., Wojno, E.D.T., Shastri, A.J., Hu, K., Cornel, L., Boedec, E., Ong, Y.-C., Chien, Y., Hunter, C.A., et al. (2011). Toxoplasma Polymorphic Effectors Determine Macrophage Polarization and Intestinal Inflammation. Cell Host & Microbe 9, 472–483. 10.1016/j.chom.2011.04.015.

25. Melo, M.B., Nguyen, Q.P., Cordeiro, C., Hassan, M.A., Yang, N., McKell, R., Rosowski, E.E., Julien, L., Butty, V., Dardé, M.-L., et al. (2013). Transcriptional Analysis of Murine Macrophages Infected with Different Toxoplasma Strains Identifies Novel Regulation of Host Signaling Pathways. PLoS Pathog 9, e1003779. 10.1371/journal.ppat.1003779.

26. Menard, K.L., Bu, L., and Denkers, E.Y. (2021). Transcriptomics analysis of Toxoplasma gondii-infected mouse macrophages reveals coding and noncoding signatures in the presence and absence of MyD88. BMC Genomics 22, 130. 10.1186/s12864-021-07437-0.

27. Pittman, K.J., Aliota, M.T., and Knoll, L.J. (2014). Dual transcriptional profiling of mice and Toxoplasma gondii during acute and chronic infection. BMC Genomics 15, 806. 10.1186/1471-2164-15-806.

28. Guha, J., Kang, B., Claudio, E., Redekar, N.R., Wang, H., Kelsall, B.L., Siebenlist, U., and Murphy, P.M. (2022). NF kappa B regulator Bcl3 controls development and function of classical dendritic cells required for resistance to Toxoplasma gondii. PLoS Pathog 18, e1010502. 10.1371/journal.ppat.1010502.

29. Avital, G., Avraham, R., Fan, A., Hashimshony, T., Hung, D.T., and Yanai, I. (2017). scDual-Seq: mapping the gene regulatory program of Salmonella infection by host and pathogen single-cell RNA-sequencing. Genome Biol 18, 200. 10.1186/s13059-017-1340-x.

30. Muñoz, J.F., Delorey, T., Ford, C.B., Li, B.Y., Thompson, D.A., Rao, R.P., and Cuomo, C.A. (2019). Coordinated host-pathogen transcriptional dynamics revealed using sorted subpopulations and single macrophages infected with Candida albicans. Nat Commun 10, 1607. 10.1038/s41467-019-09599-8.

31. Patir, A., Gossner, A., Ramachandran, P., Alves, J., Freeman, T.C., Henderson, N.C., Watson, M., and Hassan, M.A. (2020). Single-cell RNA-seq reveals CD16-monocytes as key regulators of human monocyte transcriptional response to Toxoplasma. Sci Rep 10, 21047. 10.1038/s41598-020-78250-0.

32. Sugi, T., Tomita, T., Kidaka, T., Kawai, N., Hayashida, K., Weiss, L.M., and Yamagishi, J. (2022). Single Cell Transcriptomes of In Vitro Bradyzoite Infected Cells Reveals Toxoplasma gondii Stage Dependent Host Cell Alterations. Front. Cell. Infect. Microbiol. 12, 848693. 10.3389/fcimb.2022.848693.

33. Dubey, J.P., Lindsay, D.S., and Speer, C.A. (1998). Structures of Toxoplasma gondii tachyzoites, bradyzoites, and sporozoites and biology and development of tissue cysts. Clin Microbiol Rev 11, 267–299. 10.1128/CMR.11.2.267.

34. Mosley, Y.-Y.C., and HogenEsch, H. (2017). Selection of a suitable reference gene for quantitative gene expression in mouse lymph nodes after vaccination. BMC Res Notes 10, 689. 10.1186/s13104-017-3005-y.

35. Walachowski, S., Tabouret, G., Fabre, M., and Foucras, G. (2017). Molecular Analysis of a Short-term Model of β-Glucans-Trained Immunity Highlights the Accessory Contribution of GM-CSF in Priming Mouse Macrophages Response. Front Immunol 8, 1089. 10.3389/fimmu.2017.01089.

36. McDermott, J.E., Vartanian, K.B., Mitchell, H., Stevens, S.L., Sanfilippo, A., and Stenzel-Poore, M.P. (2012). Identification and Validation of Ifit1 as an Important Innate Immune Bottleneck. PLoS ONE 7, e36465. 10.1371/journal.pone.0036465.

37. Ten Hoeve, A.L., Hakimi, M.-A., and Barragan, A. (2019). Sustained Egr-1 Response via p38 MAP Kinase Signaling Modulates Early Immune Responses of Dendritic Cells Parasitized by Toxoplasma gondii. Front Cell Infect Microbiol 9, 349. 10.3389/fcimb.2019.00349.

38. Knoll, L.J., Tomita, T., and Weiss, L.M. (2014). Chapter 15 - Bradyzoite Development. In Toxoplasma Gondii (Second Edition), L. M. Weiss and K. Kim, eds. (Academic Press), pp. 521–549. 10.1016/B978-0-12-396481-6.00015-5.

39. Plattner, F., Yarovinsky, F., Romero, S., Didry, D., Carlier, M.-F., Sher, A., and Soldati-Favre, D. (2008). Toxoplasma profilin is essential for host cell invasion and TLR11-dependent induction of an interleukin-12 response. Cell Host Microbe 3, 77–87. 10.1016/j.chom.2008.01.001.

40. Huynh, M.-H., and Carruthers, V.B. (2006). Toxoplasma MIC2 is a major determinant of invasion and virulence. PLoS Pathog 2, e84. 10.1371/journal.ppat.0020084.

41. Wang, Y., and Yin, H. (2014). Research progress on surface antigen 1 (SAG1) of Toxoplasma gondii. Parasit Vectors 7, 180. 10.1186/1756-3305-7-180.

42. Wu, R., Chen, F., Wang, N., Tang, D., and Kang, R. (2020). ACOD1 in immunometabolism and disease. Cell Mol Immunol 17, 822–833. 10.1038/s41423-020-0489-5.

43. Patin, E.C., Orr, S.J., and Schaible, U.E. (2017). Macrophage Inducible C-Type Lectin As a Multifunctional Player in Immunity. Front. Immunol. 8, 861. 10.3389/fimmu.2017.00861.

44. Zanoni, I., and Granucci, F. (2013). Role of CD14 in host protection against infections and in metabolism regulation. Front. Cell. Infect. Microbiol. 3. 10.3389/fcimb.2013.00032.

45. Zarubica, A., Trompier, D., and Chimini, G. (2007). ABCA1, from pathology to membrane function. Pflugers Arch - Eur J Physiol 453, 569–579. 10.1007/s00424-006-0108-z.

46. Gov, L., Karimzadeh, A., Ueno, N., and Lodoen, M.B. (2013). Human Innate Immunity to Toxoplasma gondii Is Mediated by Host Caspase-1 and ASC and Parasite GRA15. mBio 4, e00255–13. 10.1128/mBio.00255-13.

47. Zhao, Y., Shi, J., Shi, X., Wang, Y., Wang, F., and Shao, F. (2016). Genetic functions of the NAIP family of inflammasome receptors for bacterial ligands in mice. Journal of Experimental Medicine 213, 647–656. 10.1084/jem.20160006.

48. He, H., Brenier-Pinchart, M.-P., Braun, L., Kraut, A., Touquet, B., Couté, Y., Tardieux, I., Hakimi, M.-A., and Bougdour, A. (2018). Characterization of a Toxoplasma effector uncovers an alternative GSK3/β-catenin-regulatory pathway of inflammation. eLife 7, e39887. 10.7554/eLife.39887.

49. Jiang, A., Bloom, O., Ono, S., Cui, W., Unternaehrer, J., Jiang, S., Whitney, J.A., Connolly, J., Banchereau, J., and Mellman, I. (2007). Disruption of E-Cadherin-Mediated Adhesion Induces a Functionally Distinct Pathway of Dendritic Cell Maturation. Immunity 27, 610– 624. 10.1016/j.immuni.2007.08.015.

50. Khan, I.A., Ouellette, C., Chen, K., and Moretto, M. (2019). Toxoplasma: Immunity and Pathogenesis. Curr Clin Micro Rpt 6, 44–50. 10.1007/s40588-019-0114-5.

51. Stutz, A., Kessler, H., Kaschel, M.-E., Meissner, M., and Dalpke, A.H. (2012). Cell invasion and strain dependent induction of suppressor of cytokine signaling-1 by Toxoplasma gondii. Immunobiology 217, 28–36. 10.1016/j.imbio.2011.08.008.

52. Roy, C.N., Custodio, Á.O., de Graaf, J., Schneider, S., Akpan, I., Montross, L.K., Sanchez, M., Gaudino, A., Hentze, M.W., Andrews, N.C., et al. (2004). An Hfe-dependent pathway mediates hyposideremia in response to lipopolysaccharide-induced inflammation in mice. Nat Genet 36, 481–485. 10.1038/ng1350.

53. Thomas, R., and Yang, X. (2016). NK-DC Crosstalk in Immunity to Microbial Infection. Journal of Immunology Research 2016, 1–7. 10.1155/2016/6374379.

54. Canning, P., Ruan, Q., Schwerd, T., Hrdinka, M., Maki, J.L., Saleh, D., Suebsuwong, C., Ray, S., Brennan, P.E., Cuny, G.D., et al. (2015). Inflammatory Signaling by NOD-RIPK2 Is Inhibited by Clinically Relevant Type II Kinase Inhibitors. Chemistry & Biology 22, 1174– 1184. 10.1016/j.chembiol.2015.07.017.

55. Chan, M.M., Evans, K.W., Moore, A.R., and Fong, D. (2010). Peroxisome Proliferator-Activated Receptor (PPAR): Balance for Survival in Parasitic Infections. Journal of Biomedicine and Biotechnology 2010, 1–9. 10.1155/2010/828951.

56. Pellegrini, E., Signor, L., Singh, S., Boeri Erba, E., and Cusack, S. (2017). Structures of the inactive and active states of RIP2 kinase inform on the mechanism of activation. PLoS ONE 12, e0177161. 10.1371/journal.pone.0177161.

57. Lavine, M.D., and Arrizabalaga, G. (2009). Induction of mitotic S-phase of host and neighboring cells by Toxoplasma gondii enhances parasite invasion. Molecular and Biochemical Parasitology 164, 95–99. 10.1016/j.molbiopara.2008.11.014.

58. Molestina, R.E., El-Guendy, N., and Sinai, A.P. (2008). Infection with Toxoplasma gondii results in dysregulation of the host cell cycle. Cell Microbiol 10, 1153–1165. 10.1111/j.1462-5822.2008.01117.x.

59. van Dam, S., Võsa, U., van der Graaf, A., Franke, L., and de Magalhães, J.P. (2018). Gene co-expression analysis for functional classification and gene–disease predictions. Briefings in Bioinformatics 19, 575–592. 10.1093/bib/bbw139.

60. Drusco, A., Zanesi, N., Roldo, C., Trapasso, F., Farber, J.L., Fong, L.Y., and Croce, C.M. (2005). Knockout mice reveal a tumor suppressor function for Testin. Proc. Natl. Acad. Sci. U.S.A. 102, 10947–10951. 10.1073/pnas.0504934102.

61. Kazazian, K., Haffani, Y., Ng, D., Lee, C.M.M., Johnston, W., Kim, M., Xu, R., Pacholzyk, K., Zih, F.S.-W., Tan, J., et al. (2020). FAM46C/TENT5C functions as a tumor suppressor through inhibition of Plk4 activity. Commun Biol 3, 448. 10.1038/s42003-020-01161-3.

62. Behnke, M.S., Wootton, J.C., Lehmann, M.M., Radke, J.B., Lucas, O., Nawas, J., Sibley, L.D., and White, M.W. (2010). Coordinated Progression through Two Subtranscriptomes Underlies the Tachyzoite Cycle of Toxoplasma gondii. PLoS ONE 5, e12354. 10.1371/journal.pone.0012354.

63. Saksouk, N., Bhatti, M.M., Kieffer, S., Smith, A.T., Musset, K., Garin, J., Sullivan, W.J., Cesbron-Delauw, M.-F., and Hakimi, M.-A. (2005). Histone-Modifying Complexes Regulate Gene Expression Pertinent to the Differentiation of the Protozoan Parasite Toxoplasma gondii. Mol Cell Biol 25, 10301–10314. 10.1128/MCB.25.23.10301-10314.2005.

64. Jung, C., Lee, C.Y.-F., and Grigg, M.E. (2004). The SRS superfamily of Toxoplasma surface proteins. International Journal for Parasitology 34, 285–296. 10.1016/j.ijpara.2003.12.004.

65. Lagal, V., Binder, E.M., Huynh, M.-H., Kafsack, B.F.C., Harris, P.K., Diez, R., Chen, D., Cole, R.N., Carruthers, V.B., and Kim, K. (2010). Toxoplasma gondii protease TgSUB1 is required for cell surface processing of micronemal adhesive complexes and efficient adhesion of tachyzoites: TgSUB1 microneme protein processing. Cellular Microbiology 12, 1792–1808. 10.1111/j.1462-5822.2010.01509.x.

66. Pszenny, V., Davis, P.H., Zhou, X.W., Hunter, C.A., Carruthers, V.B., and Roos, D.S. (2012). Targeted Disruption of Toxoplasma gondii Serine Protease Inhibitor 1 Increases Bradyzoite Cyst Formation In Vitro and Parasite Tissue Burden in Mice. Infect Immun 80, 1156–1165. 10.1128/IAI.06167-11.

67. Amos, B., Aurrecoechea, C., Barba, M., Barreto, A., Basenko, E.Y., Bażant, W., Belnap, R., Blevins, A.S., Böhme, U., Brestelli, J., et al. (2022). VEuPathDB: the eukaryotic pathogen, vector and host bioinformatics resource center. Nucleic Acids Research 50, D898–D911. 10.1093/nar/gkab929.

68. Woo, Y.H., Ansari, H., Otto, T.D., Klinger, C.M., Kolisko, M., Michálek, J., Saxena, A., Shanmugam, D., Tayyrov, A., Veluchamy, A., et al. (2015). Chromerid genomes reveal the evolutionary path from photosynthetic algae to obligate intracellular parasites. Elife 4, e06974. 10.7554/eLife.06974.

69. Barylyuk, K., Koreny, L., Ke, H., Butterworth, S., Crook, O.M., Lassadi, I., Gupta, V., Tromer, E., Mourier, T., Stevens, T.J., et al. (2020). A Comprehensive Subcellular Atlas of the Toxoplasma Proteome via hyperLOPIT Provides Spatial Context for Protein Functions. Cell Host & Microbe 28, 752–766.e9. 10.1016/j.chom.2020.09.011.

70. Bradley, P.J., and Sibley, L.D. (2007). Rhoptries: an arsenal of secreted virulence factors. Current Opinion in Microbiology 10, 582–587. 10.1016/j.mib.2007.09.013.

71. Ghosh, S., Kennedy, K., Sanders, P., Matthews, K., Ralph, S.A., Counihan, N.A., and de Koning-Ward, T.F. (2017). The Plasmodium rhoptry associated protein complex is important for parasitophorous vacuole membrane structure and intraerythrocytic parasite growth. Cellular Microbiology 19, e12733. 10.1111/cmi.12733.

72. Hernández-de-los-Ríos, A., Murillo-Leon, M., Mantilla-Muriel, L.E., Arenas, A.F., Vargas-Montes, M., Cardona, N., de-la-Torre, A., Sepúlveda-Arias, J.C., and Gómez-Marín, J.E. (2019). Influence of Two Major Toxoplasma Gondii Virulence Factors (ROP16 and ROP18) on the Immune Response of Peripheral Blood Mononuclear Cells to Human Toxoplasmosis Infection. Front. Cell. Infect. Microbiol. 9, 413. 10.3389/fcimb.2019.00413.

73. Pandey, S., Gruenbaum, A., Kanashova, T., Mertins, P., Cluzel, P., and Chevrier, N. (2020). Pairwise Stimulations of Pathogen-Sensing Pathways Predict Immune Responses to Multi-adjuvant Combinations. Cell Systems 11, 495–508.e10. 10.1016/j.cels.2020.10.001.

74. Mathes, A.L., Rice, L., Affandi, A.J., DiMarzio, M., Rifkin, I.R., Stifano, G., Christmann, R.B., and Lafyatis, R. (2015). CpGB DNA activates dermal macrophages and specifically recruits inflammatory monocytes into the skin. Exp Dermatol 24, 133–139. 10.1111/exd.12603.

75. Brown, G.D. (2006). Dectin-1: a signalling non-TLR pattern-recognition receptor. Nat Rev Immunol 6, 33–43. 10.1038/nri1745.

76. Gimm, T., Wiese, M., Teschemacher, B., Deggerich, A., Schödel, J., Knaup, K.X., Hackenbeck, T., Hellerbrand, C., Amann, K., Wiesener, M.S., et al. (2010). HypoxiaLinducible protein 2 is a novel lipid droplet protein and a specific target gene of hypoxiaLinducible factorL1. FASEB j. 24, 4443–4458. 10.1096/fj.10-159806.

77. Huber, M.D., Vesely, P.W., Datta, K., and Gerace, L. (2013). Erlins restrict SREBP activation in the ER and regulate cellular cholesterol homeostasis. Journal of Cell Biology 203, 427–436. 10.1083/jcb.201305076.

78. Chiara, F., Castellaro, D., Marin, O., Petronilli, V., Brusilow, W.S., Juhaszova, M., Sollott, S.J., Forte, M., Bernardi, P., and Rasola, A. (2008). Hexokinase II Detachment from Mitochondria Triggers Apoptosis through the Permeability Transition Pore Independent of Voltage-Dependent Anion Channels. PLoS ONE 3, e1852. 10.1371/journal.pone.0001852.

79. Zhang, T., Wang, Y., Yu, H., Zhang, T., Guo, L., Xu, J., Wei, X., Wang, N., Wu, Y., Wang, X., et al. (2022). PGK1 represses autophagy-mediated cell death to promote the proliferation of liver cancer cells by phosphorylating PRAS40. Cell Death Dis 13, 68. 10.1038/s41419-022-04499-0.

80. Chen, Y., Wu, G., Li, M., Hesse, M., Ma, Y., Chen, W., Huang, H., Liu, Y., Xu, W., Tang, Y., et al. (2022). LDHA-mediated metabolic reprogramming promoted cardiomyocyte proliferation by alleviating ROS and inducing M2 macrophage polarization. Redox Biology 56, 102446. 10.1016/j.redox.2022.102446.

81. Wang, Y., Sangaré, L.O., Paredes-Santos, T.C., Hassan, M.A., Krishnamurthy, S., Furuta, A.M., Markus, B.M., Lourido, S., and Saeij, J.P.J. (2020). Genome-wide screens identify Toxoplasma gondii determinants of parasite fitness in IFNγ-activated murine macrophages. Nat Commun 11, 5258. 10.1038/s41467-020-18991-8.

82. Jeidane, S., Scott-Boyer, M.-P., Tremblay, N., Cardin, S., Picard, S., Baril, M., Lamarre, D., and Deschepper, C.F. (2016). Association of a Network of Interferon-Stimulated Genes with a Locus Encoding a Negative Regulator of Non-conventional IKK Kinases and IFNB1. Cell Reports 17, 425–435. 10.1016/j.celrep.2016.09.009.

83. Lou, Y., Han, M., Liu, H., Niu, Y., Liang, Y., Guo, J., Zhang, W., and Wang, H. (2020). Essential roles of S100A10 in Toll-like receptor signaling and immunity to infection. Cell Mol Immunol 17, 1053–1062. 10.1038/s41423-019-0278-1.

84. Zou, B., Liu, J., Klionsky, D.J., Tang, D., and Kang, R. (2020). Extracellular SQSTM1 as an inflammatory mediator. Autophagy 16, 2313–2315. 10.1080/15548627.2020.1843253.

85. Pooja, S., Pushpanathan, M., Gunasekaran, P., and Rajendhran, J. (2015). EndocytosisLMediated Invasion and Pathogenicity of Streptococcus agalactiae in Rat Cardiomyocyte (H9C2). PLoS ONE 10, e0139733. 10.1371/journal.pone.0139733.

86. Case, N.T., Duah, K., Larsen, B., Wong, C.J., Gingras, A.-C., O’Meara, T.R., Robbins, N., Veri, A.O., Whitesell, L., and Cowen, L.E. (2021). The macrophage-derived protein PTMA induces filamentation of the human fungal pathogen Candida albicans. Cell Reports 36, 109584. 10.1016/j.celrep.2021.109584.

87. Domogalla, M.P., Rostan, P.V., Raker, V.K., and Steinbrink, K. (2017). Tolerance through Education: How Tolerogenic Dendritic Cells Shape Immunity. Front. Immunol. 8, 1764. 10.3389/fimmu.2017.01764.

88. Manicassamy, S., and Pulendran, B. (2011). Dendritic cell control of tolerogenic responses. Immunol Rev 241, 206–227. 10.1111/j.1600-065X.2011.01015.x.

89. Bhattacharya, P., Thiruppathi, M., Elshabrawy, H.A., Alharshawi, K., Kumar, P., and Prabhakar, B.S. (2015). GM-CSF: An immune modulatory cytokine that can suppress autoimmunity. Cytokine 75, 261–271. 10.1016/j.cyto.2015.05.030.

90. Schutt, C.R., Gendelman, H.E., and Mosley, R.L. (2018). Tolerogenic bone marrow-derived dendritic cells induce neuroprotective regulatory T cells in a model of Parkinson’s disease. Mol Neurodegeneration 13, 26. 10.1186/s13024-018-0255-7.

91. Steichen, A.L., Binstock, B.J., Mishra, B.B., and Sharma, J. (2013). C-type lectin receptor Clec4d plays a protective role in resolution of Gram-negative pneumonia. J Leukoc Biol 94, 393–398. 10.1189/jlb.1212622.

92. Yonekawa, A., Saijo, S., Hoshino, Y., Miyake, Y., Ishikawa, E., Suzukawa, M., Inoue, H., Tanaka, M., Yoneyama, M., Oh-hora, M., et al. (2014). Dectin-2 Is a Direct Receptor for Mannose-Capped Lipoarabinomannan of Mycobacteria. Immunity 41, 402–413. 10.1016/j.immuni.2014.08.005.

93. Jeong, Y.-I., Hong, S.-H., Cho, S.-H., Park, M.Y., and Lee, S.-E. (2016). Induction of IL-10-producing regulatory B cells following Toxoplasma gondii infection is important to the cyst formation. Biochem Biophys Rep 7, 91–97. 10.1016/j.bbrep.2016.05.008.

94. Sullivan, W.J., and Jeffers, V. (2012). Mechanisms of Toxoplasma gondii persistence and latency. FEMS Microbiol Rev 36, 717–733. 10.1111/j.1574-6976.2011.00305.x.

95. Pick, R., Begandt, D., Stocker, T.J., Salvermoser, M., Thome, S., Böttcher, R.T., Montanez, E., Harrison, U., Forné, I., Khandoga, A.G., et al. (2017). Coronin 1A, a novel player in integrin biology, controls neutrophil trafficking in innate immunity. Blood 130, 847– 858. 10.1182/blood-2016-11-749622.

96. Tinoco, R., Neubert, E.N., Stairiker, C.J., Henriquez, M.L., and Bradley, L.M. (2021). PSGL-1 Is a T Cell Intrinsic Inhibitor That Regulates Effector and Memory Differentiation and Responses During Viral Infection. Front Immunol 12, 677824. 10.3389/fimmu.2021.677824.

97. Gannoun-Zaki, L., Pätzold, L., Huc-Brandt, S., Baronian, G., Elhawy, M.I., Gaupp, R., Martin, M., Blanc-Potard, A.-B., Letourneur, F., Bischoff, M., et al. (2018). PtpA, a secreted tyrosine phosphatase from Staphylococcus aureus, contributes to virulence and interacts with coronin-1A during infection. J Biol Chem 293, 15569–15580. 10.1074/jbc.RA118.003555.

98. Yu, Z., Lu, Y., Liu, Z., Aleem, M.T., Liu, J., Luo, J., Yan, R., Xu, L., Song, X., and Li, X. (2021). Recombinant Toxoplasma gondii Ribosomal Protein P2 Modulates the Functions of Murine Macrophages In Vitro and Provides Immunity against Acute Toxoplasmosis In Vivo. Vaccines 9, 357. 10.3390/vaccines9040357.

99. Cérède, O., Dubremetz, J.F., Soête, M., Deslée, D., Vial, H., Bout, D., and Lebrun, M. (2005). Synergistic role of micronemal proteins in Toxoplasma gondii virulence. J Exp Med 201, 453–463. 10.1084/jem.20041672.

100. Mercier, C., Howe, D.K., Mordue, D., Lingnau, M., and Sibley, L.D. (1998). Targeted disruption of the GRA2 locus in Toxoplasma gondii decreases acute virulence in mice. Infect Immun 66, 4176–4182. 10.1128/IAI.66.9.4176-4182.1998.

101. Lebrun, M., Carruthers, V.B., and Cesbron-Delauw, M.-F. (2014). Toxoplasma Secretory Proteins and Their Roles in Cell Invasion and Intracellular Survival. In Toxoplasma Gondii (Elsevier), pp. 389–453. 10.1016/B978-0-12-396481-6.00012-X.

102. Butcher, B.A., Fox, B.A., Rommereim, L.M., Kim, S.G., Maurer, K.J., Yarovinsky, F., Herbert, D.R., Bzik, D.J., and Denkers, E.Y. (2011). Toxoplasma gondii Rhoptry Kinase ROP16 Activates STAT3 and STAT6 Resulting in Cytokine Inhibition and Arginase-1-Dependent Growth Control. PLoS Pathog 7, e1002236. 10.1371/journal.ppat.1002236.

103. Song, J.S., Kim, Y.-J., Han, K.U., Yoon, B.D., and Kim, J.W. (2015). Zymosan and PMA activate the immune responses of Mutz3-derived dendritic cells synergistically. Immunology Letters 167, 41–46. 10.1016/j.imlet.2015.07.002.

104. Hartmann, G. (2003). CpG: unraveling the key to B-cell function. Blood 101, 4230–4231. 10.1182/blood-2003-03-0921.

105. He, B., Qiao, X., and Cerutti, A. (2004). CpG DNA Induces IgG Class Switch DNA Recombination by Activating Human B Cells through an Innate Pathway That Requires TLR9 and Cooperates with IL-10. J Immunol 173, 4479–4491. 10.4049/jimmunol.173.7.4479.

106. Yarovinsky, F., and Sher, A. (2006). Toll-like receptor recognition of Toxoplasma gondii. Int J Parasitol 36, 255–259. 10.1016/j.ijpara.2005.12.003.

107. Khan, I.A. (2007). Toll road for Toxoplasma gondii: the mystery continues. Trends in Parasitology 23, 1–3. 10.1016/j.pt.2006.10.001.

108. Mun, H.-S. (2003). TLR2 as an essential molecule for protective immunity against Toxoplasma gondii infection. International Immunology 15, 1081–1087. 10.1093/intimm/dxg108.

109. Kanatani, S., Fuks, J.M., Olafsson, E.B., Westermark, L., Chambers, B., Varas-Godoy, M., Uhlén, P., and Barragan, A. (2017). Voltage-dependent calcium channel signaling mediates GABAA receptor-induced migratory activation of dendritic cells infected by Toxoplasma gondii. PLoS Pathog 13, e1006739. 10.1371/journal.ppat.1006739.

110. Fuks, J.M., Arrighi, R.B.G., Weidner, J.M., Kumar Mendu, S., Jin, Z., Wallin, R.P.A., Rethi, B., Birnir, B., and Barragan, A. (2012). GABAergic Signaling Is Linked to a Hypermigratory Phenotype in Dendritic Cells Infected by Toxoplasma gondii. PLoS Pathog 8, e1003051. 10.1371/journal.ppat.1003051.

111. Norbury, C.C., Chambers, B.J., Prescott, A.R., Ljunggren, H.-G., and Watts, C. (1997). Constitutive macropinocytosis allows TAP-dependent major histocompatibility compex class I presentation of exogenous soluble antigen by bone marrow-derived dendritic cells. Eur. J. Immunol. 27, 280–288. 10.1002/eji.1830270141.

112. Hitziger, N., Dellacasa, I., Albiger, B., and Barragan, A. (2005). Dissemination of Toxoplasma gondii to immunoprivileged organs and role of Toll/interleukin-1 receptor signalling for host resistance assessed by in vivo bioluminescence imaging. Cell Microbiol 7, 837–848. 10.1111/j.1462-5822.2005.00517.x.

113. Rodrigues, O.R., and Monard, S. (2016). A rapid method to verify single-cell deposition setup for cell sorters. Cytometry A 89, 594–600. 10.1002/cyto.a.22865.

114. Picelli, S., Björklund, å.K., Faridani, O.R., Sagasser, S., Winberg, G., and Sandberg, R. (2013). Smart-seq2 for sensitive full-length transcriptome profiling in single cells. Nature Methods 10, 1096–1098. 10.1038/nmeth.2639.

115. Stuart, T., Butler, A., Hoffman, P., Hafemeister, C., Papalexi, E., Mauck, W.M., Hao, Y., Stoeckius, M., Smibert, P., and Satija, R. (2019). Comprehensive Integration of Single-Cell Data. Cell 177, 1888–1902.e21. 10.1016/j.cell.2019.05.031.

116. Hafemeister, C., and Satija, R. (2019). Normalization and variance stabilization of single-cell RNA-seq data using regularized negative binomial regression. Genome Biol 20, 296. 10.1186/s13059-019-1874-1.

117. Fu, R., Gillen, A.E., Sheridan, R.M., Tian, C., Daya, M., Hao, Y., Hesselberth, J.R., and Riemondy, K.A. (2020). clustifyr: an R package for automated single-cell RNA sequencing cluster classification. F1000Res 9, 223. 10.12688/f1000research.22969.2.

118. Tirosh, I., Izar, B., Prakadan, S.M., Wadsworth, M.H., Treacy, D., Trombetta, J.J., Rotem, A., Rodman, C., Lian, C., Murphy, G., et al. (2016). Dissecting the multicellular ecosystem of metastatic melanoma by single-cell RNA-seq. Science 352, 189–196. 10.1126/science.aad0501.

119. Camon, E.B., Barrell, D.G., Dimmer, E.C., Lee, V., Magrane, M., Maslen, J., Binns, D., and Apweiler, R. (2005). An evaluation of GO annotation retrieval for BioCreAtIvE and GOA. BMC Bioinformatics 6 Suppl 1, S17. 10.1186/1471-2105-6-S1-S17.

120. Love, M.I., Huber, W., and Anders, S. (2014). Moderated estimation of fold change and dispersion for RNA-seq data with DESeq2. Genome Biol 15, 550. 10.1186/s13059-014-0550-8.

