## Supplementary figures for "scDual-Seq of *Toxoplasma gondii*-infected mouse bone marrow-derived dendritic cells reveals host cell heterogeneity and differential infection dynamics"

**Supplementary figure 1: Percentage of reads per cell mapped to the genome of *M.musculus* or *T. gondii* (ME49).** Dark color (blue) indicates mouse as species, light color (orange) indicates *T. gondii* (left panel). Distribution of unique *T. gondii* specific molecules across infection conditions (strain and time point) in percent per cell (right panel).

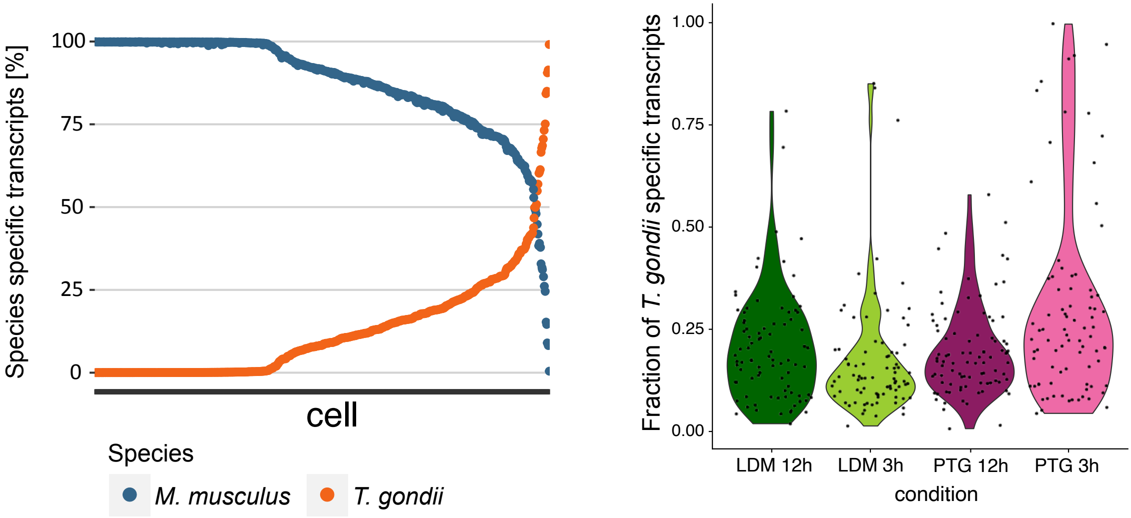

**Supplementary figure 2: Distribution of original infection conditions across identified clusters in murine data (M1-M9).** For each cluster (M1-M9) the percentage of cells belonging to the original condition is shown in a stacked barplot from 0 to 100%. Each color signifies a different condition depicted on the legend (right).

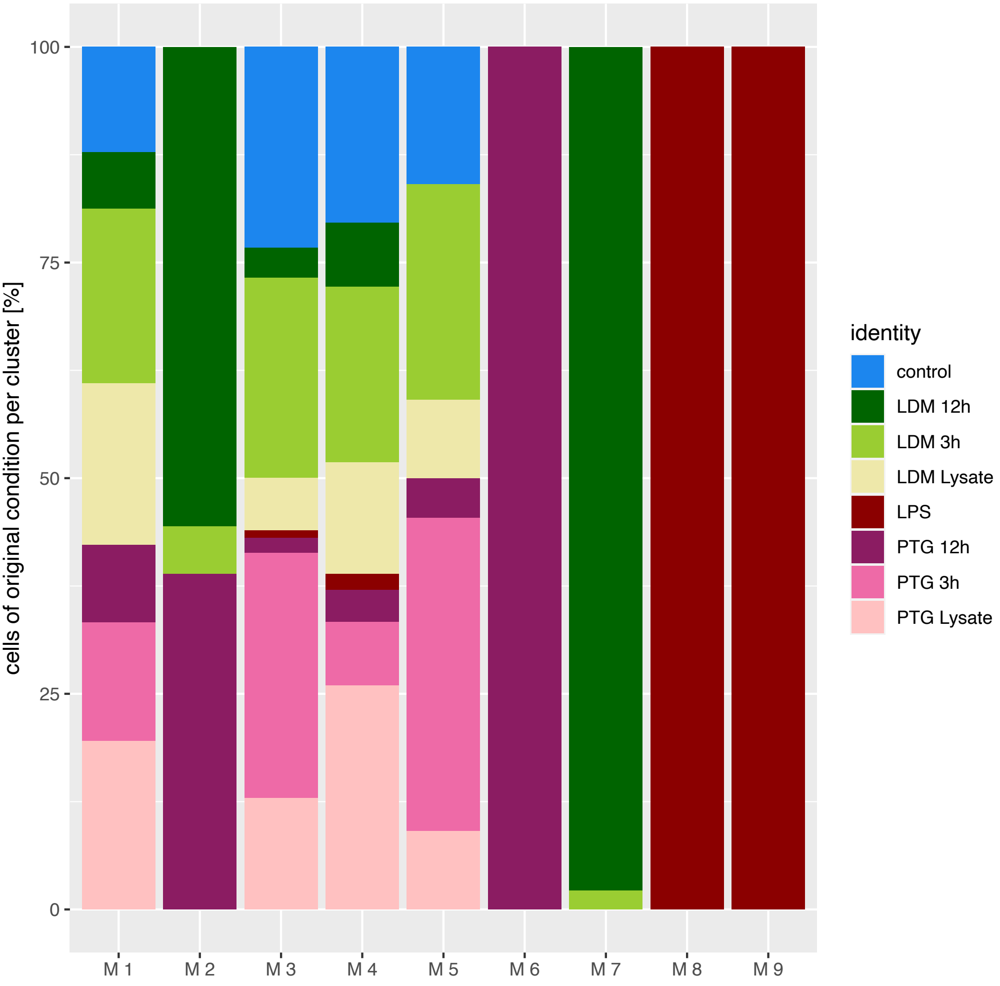

**Supplementary figure 3: Gene expression signature of identified BMDC subpopulations.** Heatmap depicting averaged gene expression profiles of cells identified as GM-DC and GM-Mac across murine clusters M1-M9. Genes identified as markers for each cell type as well as CD surface marker genes are highlighted with a black line. High expression is shown in yellow, while low expression is shown in dark purple.

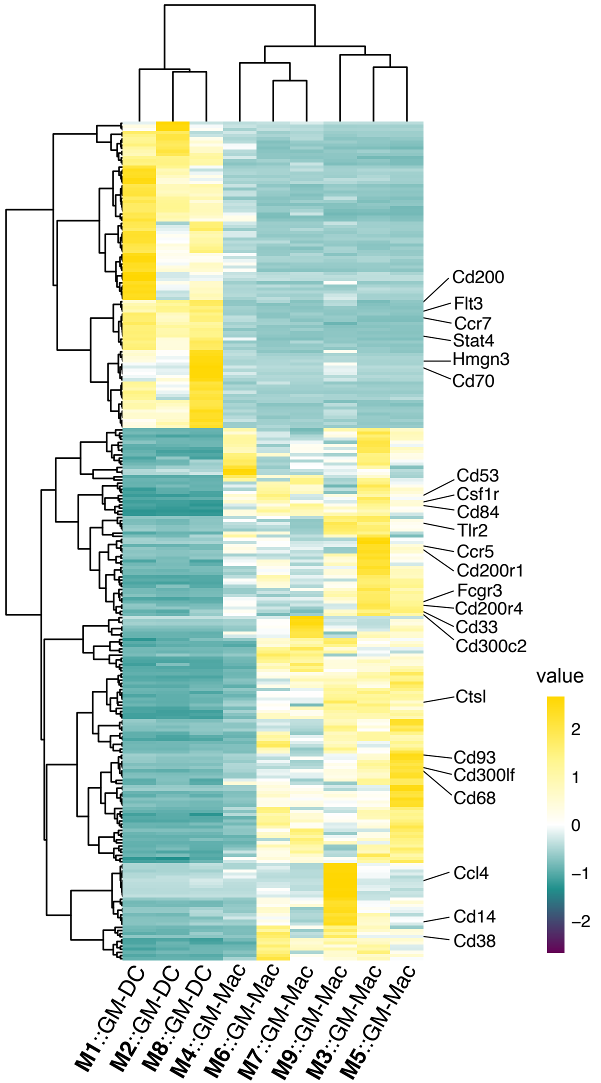

**Supplementary figure 4: Cell cycle scoring of BMDCs infected with *T. gondii* parasites.** UMAP projection of cells grouped by assigned cell cycle phase using the *CellCycleScoring* function in Seurat (methods).

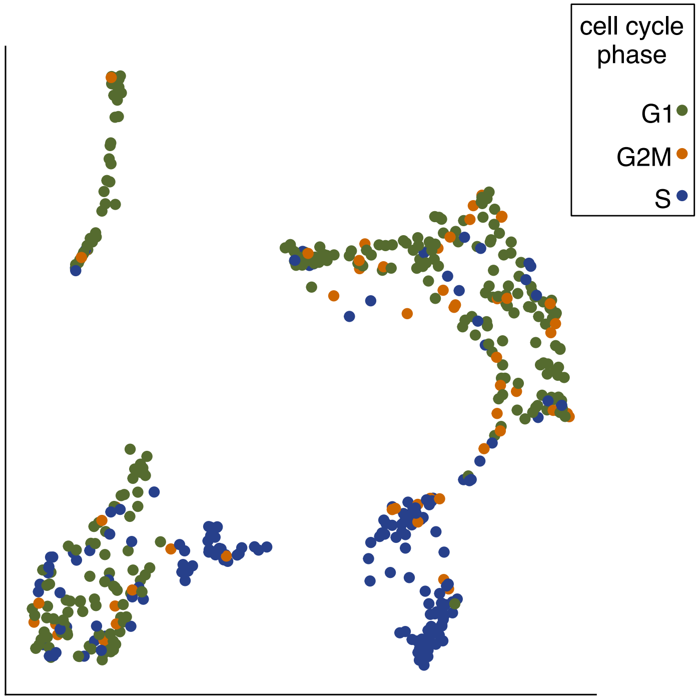

**Supplementary figure 5: Expression profiles of selected genes encoding cell cycle regulation genes (cyclins and kinases). a** Density plot exhibiting expression values of gene expression for *Ccnd2* (left), *Ccne1* and *Ccne2* encoding CyclinD2, CyclinE1 and CyclinE2 across murine clusters. Density distributions are colored based on the cluster identity as seen in the legend (right). **b** Density plot exhibiting expression values of gene expression for Cdk1 (left), Cdk2 (left middle), Cdk4 (right middle) and Cdk6 (right) encoding cyclin depending kinases corresponding phosphorilating cyclins during cell cycle progression. Density distributions are colored based on the cluster identity as seen in the legend (right).

**a**

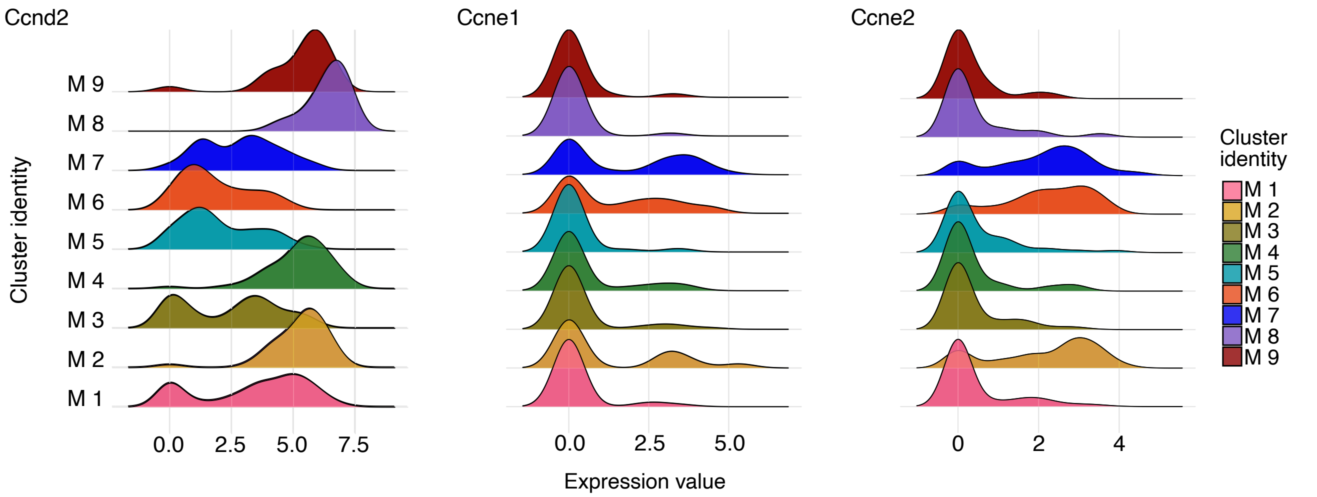


**b**

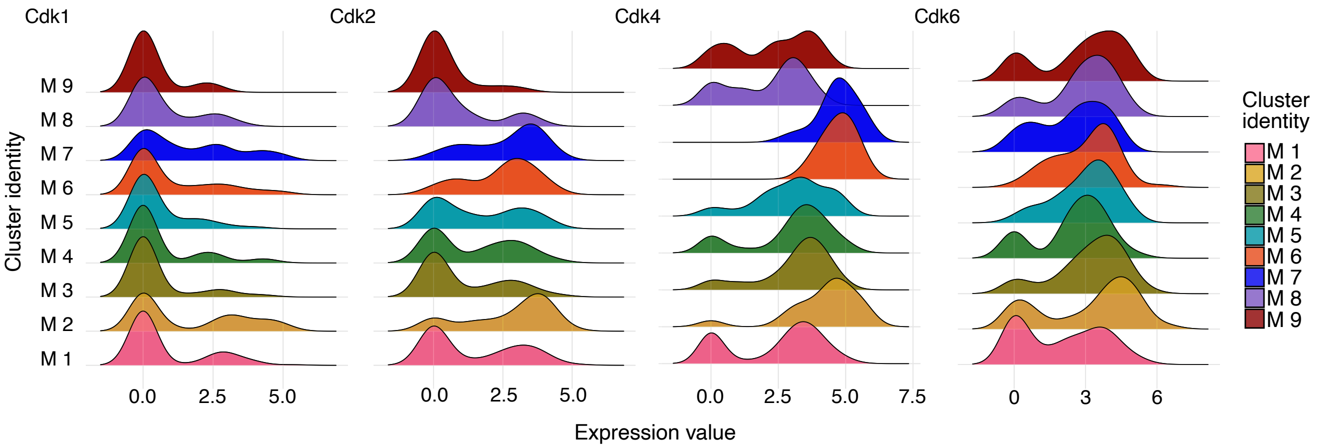

**Supplementary figure 6: Co-expression network analysis of differentially expressed genes between murine clusters M6 (PTG, 12 hpi) and M7 (LDM, 12 hpi).** Correlation based co-expression analysis (see methods for details) of marker genes of murine clusters M6 and M7. Each node represents a gene, with the weight of each gene illustrated by the size of the node. The color of the node specifies the origin or species of the marker gene, with M7 marker genes represented in turquoise and M6 marker genes in orange. Grey nodes represent murine co-expressed genes while yellow nodes represent *T. gondii* co-expressed genes. The directionality of the arrow indicates the direction of the correlation. The width of the arrow indicates the correlation value with increasing width upon increasing values. Several connected nodes represent one interaction cluster. Interaction clusters shown in figure 2b are highlighted with a turquoise and orange frame, respectively.

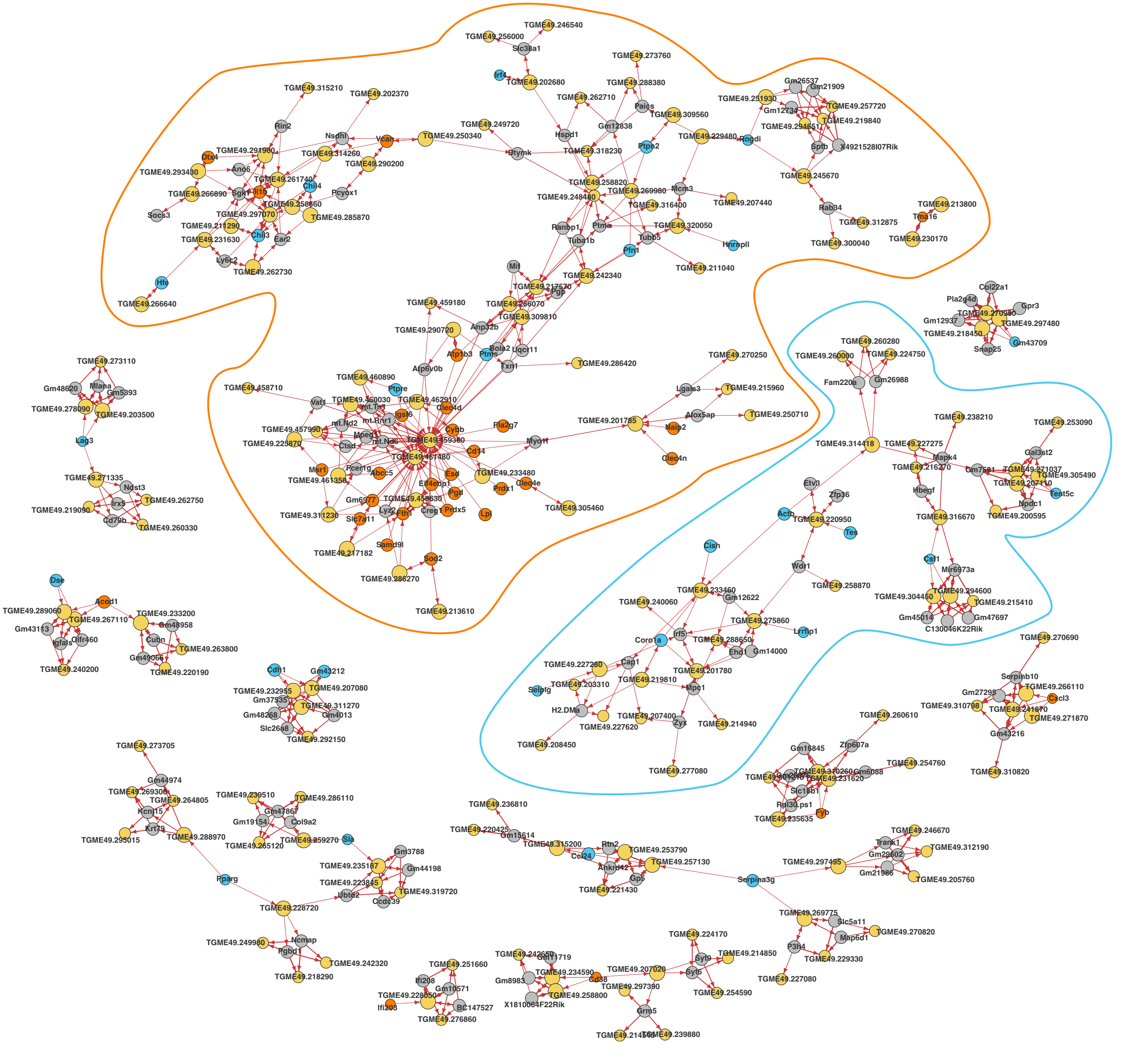

**Supplementary figure 7: Correlated genes of extended *T. gondii* network 1 (ETGN1).** Visualization of Pearson correlation coefficients between genes in extended *T. gondii* ETGN 1. Correlation values are shown in a gradient from most negative values in dark purple to most positive values in yellow (light).

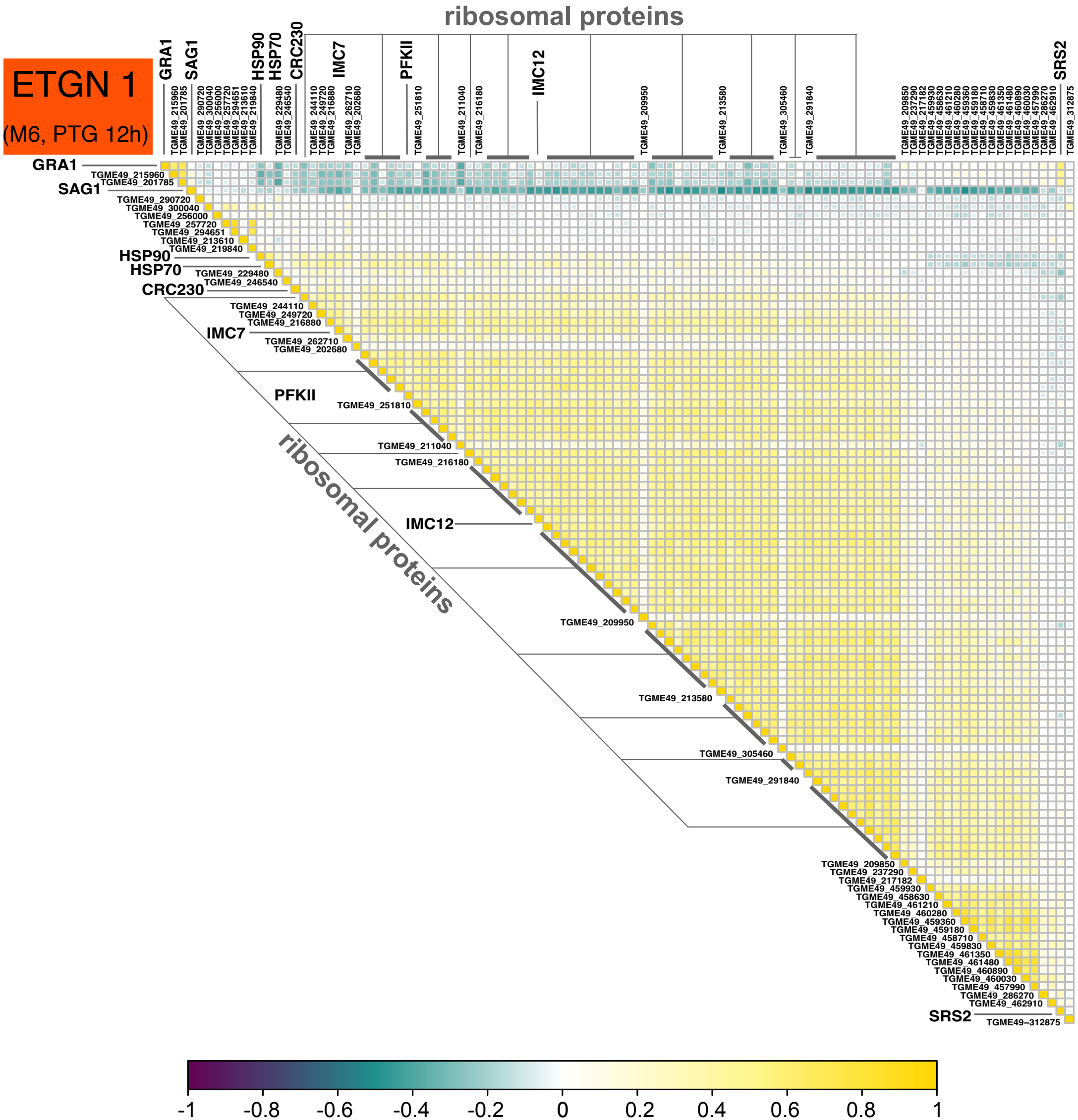

**Supplementary figure 8: Correlated genes of extended *T. gondii* network 2 (ETGN2).** Visualization of Pearson correlation coefficients between genes in extended *T. gondii* ETGN 2. Correlation values are shown in a gradient from most negative values in dark purple to most positive values in yellow (light).

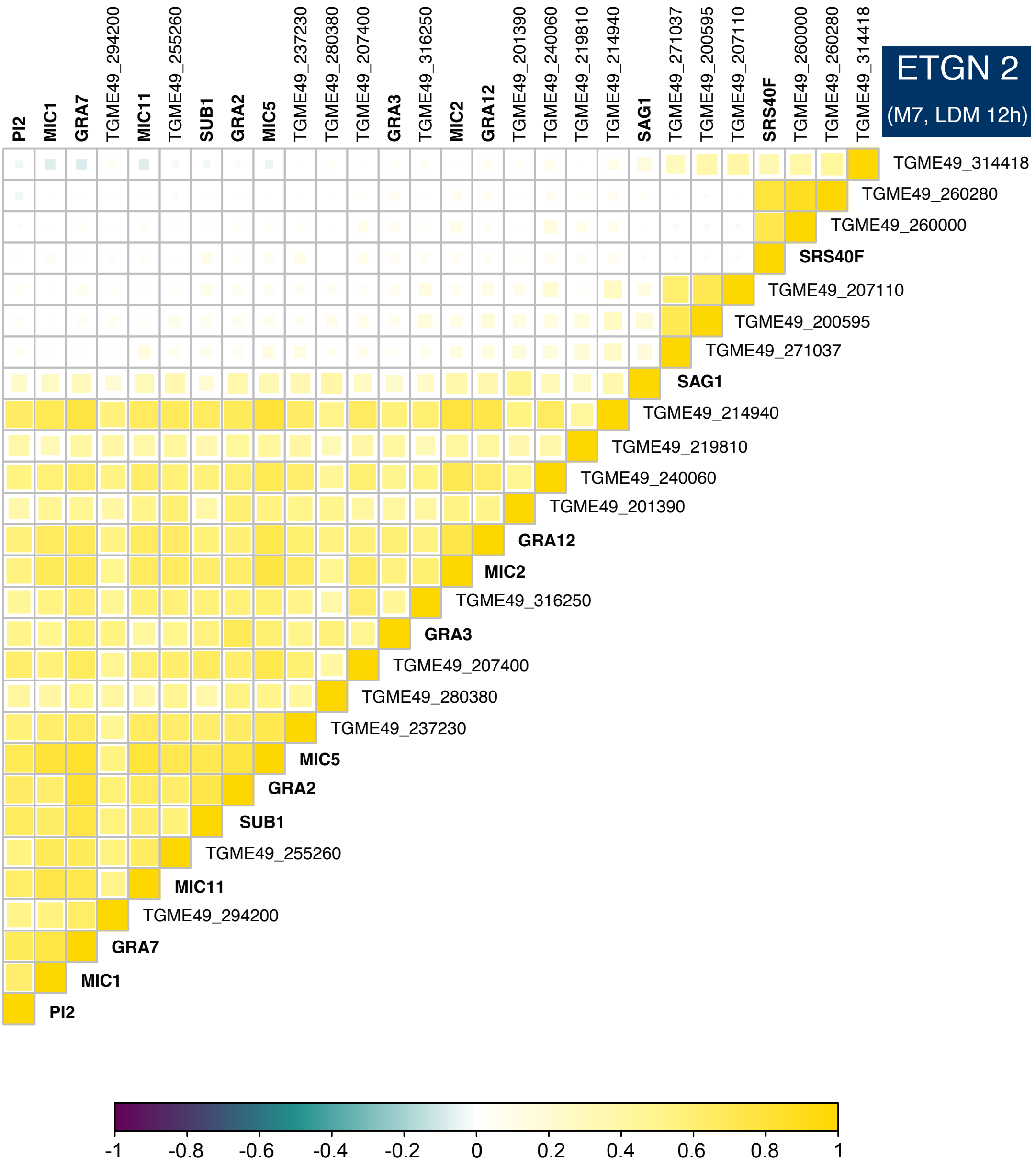

**Supplementary figure 9: Effect of cell cycle regression on clustering of *T. gondii* parasites.** UMAP projection of cells grouped by assigned replication state of the parasite using a modified version of the CellCycleScoring function in Seurat and known G1 and S/M transcription profiles of the parasite (methods) before regression of the assigned cell cycle state (left panel). UMAP projection of cells grouped by assigned replication state of the parasite using a modified version of the CellCycleScoring function in Seurat and known G1 and S/M transcription profiles of the parasite (methods) after regression of the assigned cell cycle state (right panel).

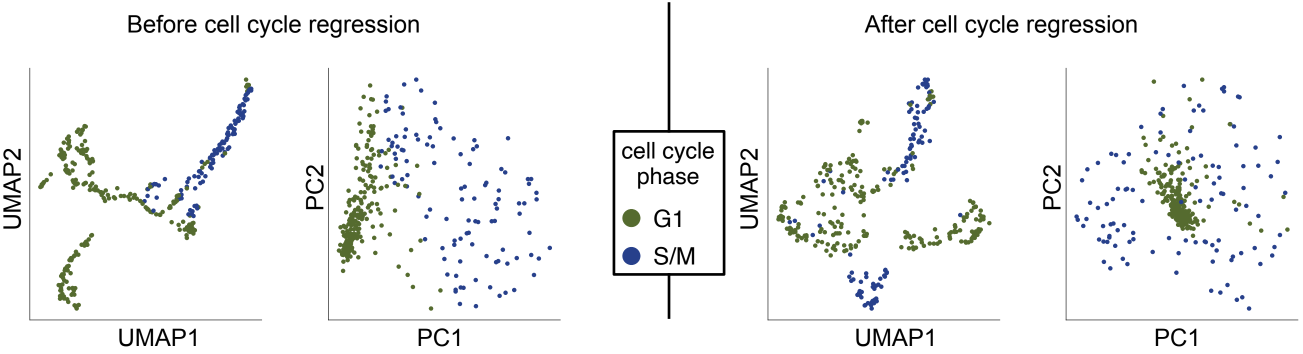

**Supplementary figure 10: Differentially expressed genes between identified *T. gondii* clusters.** Heatmap depicting averaged gene expression of *T. gondii* marker genes of identified clusters. High expression is shown in yellow, while low expression is shown in dark purple.

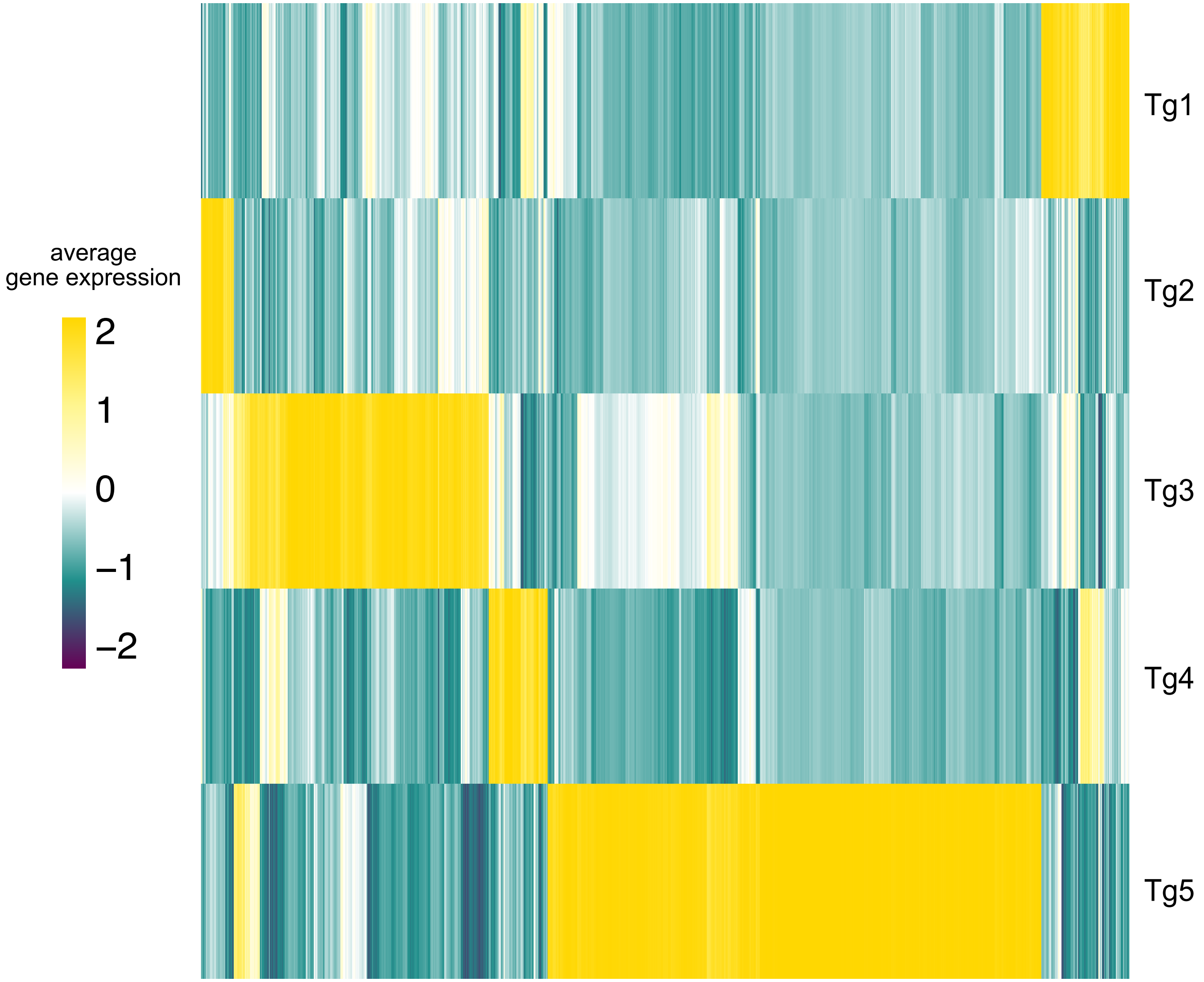

**Supplementary figure 11: Heatmap depicting differentially expressed *T. gondii* genes grouped by *T. gondii* clusters (Tg1- Tg4) across host clusters exhibiting infection (M1-M7).** Cell type annotations of host clusters are indicated by turquoise (GM-DC) and purple (GM-Mac). Averaged gene expression is shown in a color gradient from low expression (dark purple) to high expression (yellow). The *T. gondii* clusters are represented in the colors selected for each cluster

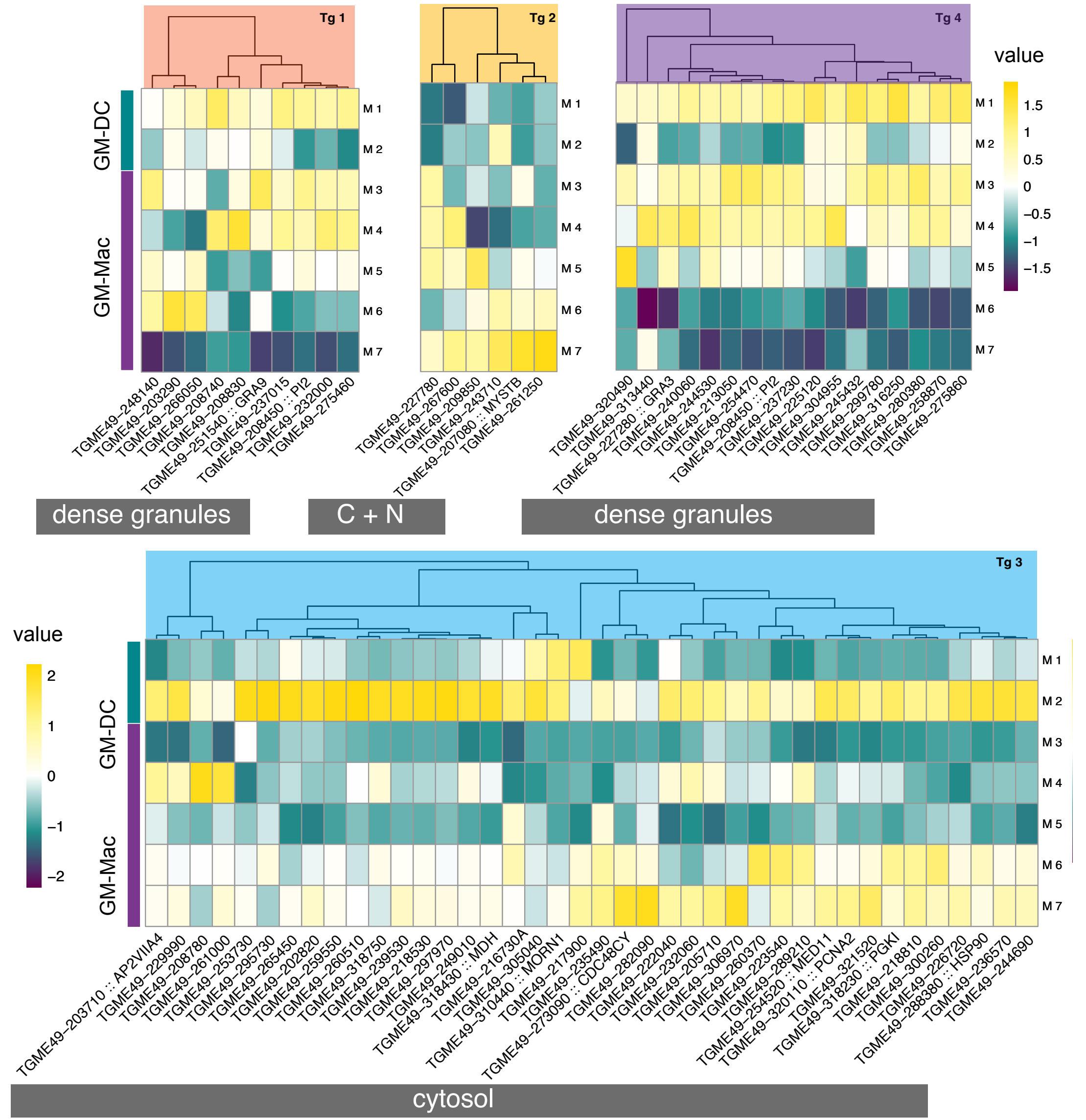

**Supplementary figure 12: Heatmap depicting differentially expressed *T. gondii* genes of *T. gondii* cluster Tg5 across host clusters exhibiting infection (M1-M7).** Cell type annotations of host clusters are indicated by turquoise (GM-DC) and purple (GM-Mac). Averaged gene expression is shown in a color gradient from low expression (dark purple) to high expression (yellow). The *T. gondii* clusters are represented in the colors selected for each cluster

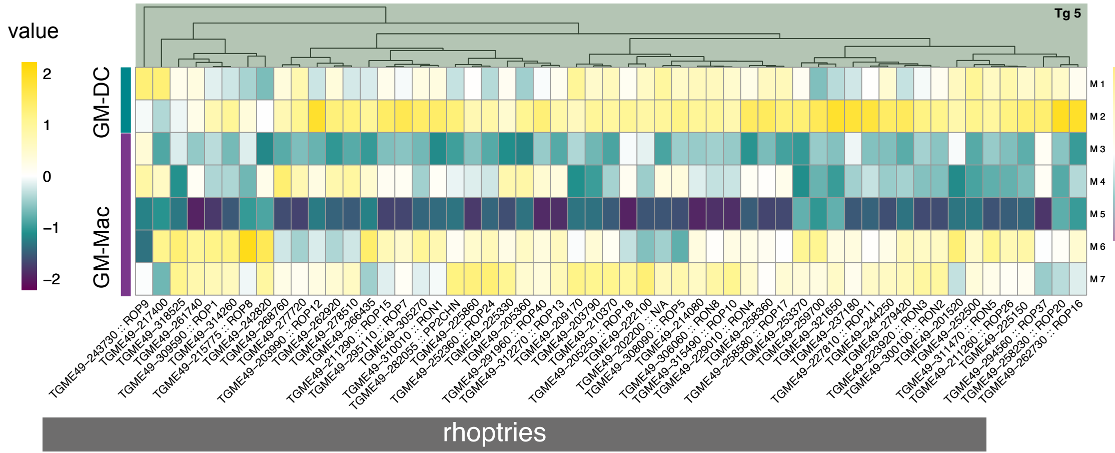

**Supplementary figure 13: Co-expression network analysis for genes exhibiting correlation with gene expression of other host and *T. gondii* genes.** Input seed genes shown to be essential under INF-stress **(Wang et al., 2020)** and present in our data are shown in yellow. Correlating host genes are depicted as purple nodes while correlating *T. gondii* genes are shown as gray nodes. The directionality of the arrow indicates the direction of the interaction of a gene and its interactor and the width of the arrow indicates the strength of the interaction. The red color of the edges indicate only positive interactions occur using our settings and several connected nodes represent one interaction cluster. The two largest co-expression clusters, depicted in main figure 5 are shown on a grey background.

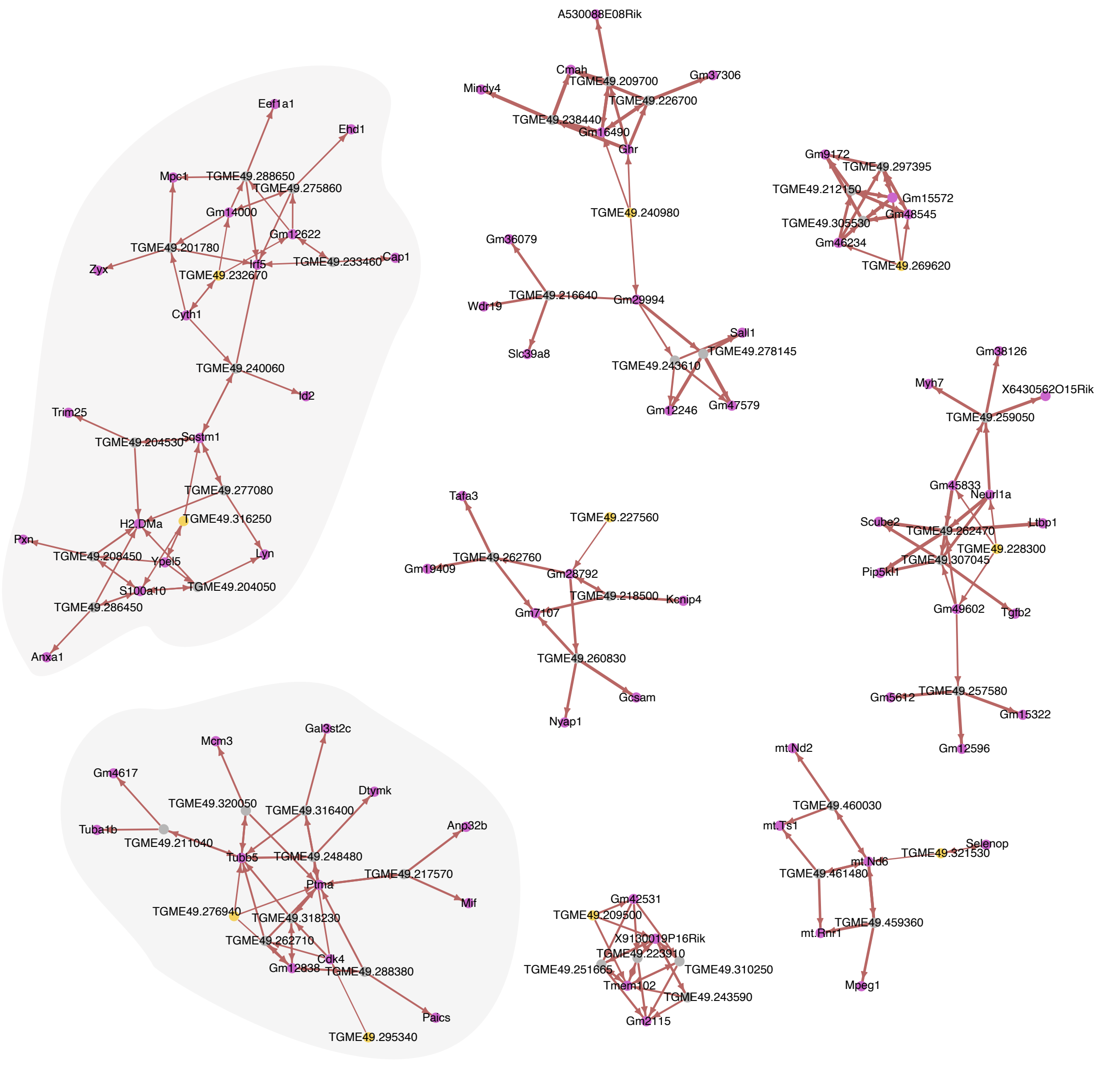
